# Task and Behavior-Related Variables Are Encoded by the Postrhinal and Medial Entorhinal Cortex During Non-Spatial Associative Learning

**DOI:** 10.1101/2024.12.20.629630

**Authors:** Ingeborg Nymoen Nystuen, Frederik Sebastian Rogge, Anna Hjertvik Aasen, Sverre Grødem, Anders Malthe-Sørenssen, Mikkel Elle Lepperød, Torkel Hafting, Marianne Fyhn, Kristian Kinden Lensjø

**Affiliations:** Department of Bioscience, University of Oslo, Norway; Institute of Basic Medical Sciences, University of Oslo, Norway; Department of Physics, University of Oslo, Norway

## Abstract

The medial entorhinal cortex (MEC) is pivotal in spatial computations and episodic memory. In particular, an animal’s position can be decoded from the activity of entorhinal grid cells. However, it remains elusive whether MEC could play a more general role in different types of associative learning and how the representations develop during the learning process. It has been shown that the postrhinal cortex (POR), which is directly connected to MEC, integrates visual stimuli with salient outcomes. Here, we use a non-spatial visual association task to investigate whether MEC neurons represent low-level visual cues during learning. Using a Go/NoGo visual association task, we recorded neural activity in MEC and POR throughout the learning phase as mice associated drifting gratings with rewarded, aversive, or neutral outcomes.

Our findings reveal that the neural tuning curves in both the POR and MEC change with the learning of the task. From the start of training, the POR neurons exhibited response tuning to the visual cues, and the tuning was stable to cue orientations during learning. In contrast, MEC neurons did not initially respond very strongly to visual cues but developed a robust tuning toward the rewarded trials. While the MEC’s representation of visual information was limited, it encoded other task elements. A large fraction of the neurons formed distinct functional clusters that were either activated or suppressed by reward-related behavior. Remarkably, these clusters segregated anatomically in MEC and maintained strong within-cluster correlations before and after training. Notably, although the same functional clusters were apparent in the POR, they did not show any anatomical structure as in the MEC. Task reversal induced significant changes in network responses across both regions, with a decrease in overall task-responsive neurons but a slight increase in stimulus representation. Strikingly, information about the choice to lick emerged with learning in both brain areas, and most significantly within the functional cell clusters representing reward consumption and plus-cue stimulus. Our results demonstrate that although neurons in MEC and POR develop behavior-modulated tuning during learning of a non-spatial visual association task, the MEC exhibits stronger within-cluster correlations and anatomical organization. Conversely, the POR population exhibits less structural organization and more specific stimulus-tuning, which is reflective of being a higher visual association area. Our findings reveal that the MEC can encode task– and behavior-related variables beyond spatial information.

## 1 Introduction

A fundamental function of the brain is to assign meaning to encountered stimuli to make appropriate decisions and actions. Forming associations between sensory cues and salient outcomes necessitates flexible neural encoding, which allows for the decoding of stimuli and evaluating their valence in a constantly changing environment. Even with familiar stimuli, the appropriate responses and expected outcomes can vary depending on context and the relevant rules of engagement [Ramesh et al., 2018, McGuire et al., 2022]. Representations of stimulus-outcome associations have been observed across a wide range of brain regions, with neural correlates of stimulus, choice, and outcome localized to specific cortical and subcortical areas [Steinmetz et al., 2019, Lutas et al., 2019]. Notably, modality-specific representations of stimulus-outcome associations have been causally linked to areas of the parahippocampal region [Lee et al., 2021, Ramesh et al., 2018].

The postrhinal cortex (POR) is one of the integration sites of the lateral visual association cortex, linking visual stimuli with salient outcomes [Burgess et al., 2016, Ramesh et al., 2018, Sugden et al., 2020]. In POR, the reward bias that develops with cue-outcome associations is regulated by motivation-dependent input from the lateral amygdala [Burgess et al., 2016]. In addition to its role in non-spatial associations, the POR also integrates visual information within spatial contexts, potentially providing visuospatial input to the medial entorhinal cortex (MEC) [LaChance et al., 2022, Koganezawa et al., 2015] but see [Doan et al., 2019, Nilssen et al., 2019].

The MEC, situated downstream of the POR, has been extensively studied for its crucial role in spatial computations. It contains a variety of functional cell types that correspond to specific behaviors and spatial features of the environment [Sargolini et al., 2006, Kropff et al., 2015, Solstad et al., 2008, Fyhn et al., 2004, Hafting et al., 2005], and is thought to provide the hippocampus with spatial anchors in episodic memory processing [Mulders et al., 2021]. Notably, recent experiments from mice navigating virtual reality have revealed that MEC cell subpopulations are tuned to multimodal (auditory and visual) and unimodal cues [Nguyen et al., 2024]. These findings align with other studies showing that MEC cells not only generally anchor to visually distinguishable landmarks [Kinkhabwala et al., 2020, Campbell et al., 2018] but also represent sound sequences in an auditory task ([Aronov et al., 2017]). Additionally, cells in the MEC have been linked to non-spatial functions evident in cells encoding temporal aspects of behavioral tasks [Kraus et al., 2015, Heys and Dombeck, 2018] and reward representations, with grid cell structures reorganizing around goal locations [Boccara et al., 2019, Butler et al., 2019]. Although the origin of reward modulation signals to the MEC remains elusive, anatomical evidence indicates that the MEC is innervated by the amygdala, both directly [Krettek and Price, 1974, Pitkänen et al., 2002] and through POR originating neurons [Meier et al., 2021].

Given the connectivity between the regions, the well-mapped representations of visual associations in POR [Burgess et al., 2016, Ramesh et al., 2018, Sugden et al., 2020, McGuire et al., 2022], and emerging evidence on MEC function in stimulus encoding in spatial tasks Nguyen et al. [2024], Kinkhabwala et al. [2020], we hypothesized that the MEC would encode rewarded stimulus-outcome associations also in a non-spatial context. Specifically, we aimed to investigate whether MEC neurons represent low-level visual cues and how neural populations in the MEC respond to outcome-based association learning compared to the upstream POR.

To address this, we recorded neural activity throughout the learning phase, from naïve to expert performance, as mice learned to associate drifting gratings with rewarding, aversive, or neutral outcomes in a visual Go/NoGo task. By tracking the same population across sessions, we observed that a substantial fraction of POR and MEC neurons exhibited dynamic changes in their response profiles as learning progressed. Consistent with previous reports, POR neurons showed stable tuning to cue orientations, with a bias toward the rewarded cue as mice successfully paired cues and outcomes. Strikingly, MEC neurons exhibited a strong bias toward rewarded trials, with some neurons maintaining sensitivity to cue identity even after cue-outcome reversals. Although the visual information alone represented by MEC cells was limited, the behavioral approach, which in our task takes the form of anticipatory licking behavior, was represented by a significant fraction of the population, with clear anatomical separation between the representations. Indeed, we find that information about the behavioral choice is represented in these neurons prior to licking, a feature that is absent in naive mice but develops with learning. Our findings demonstrate that behavior-modulated cells follow similar developmental patterns across both regions; however, the MEC’s correlation structure and anatomical organization are more pronounced. In contrast, the POR exhibits less structural organization but more specific stimulus tuning, which is limited in the MEC.

## 2 Results

### Mice successfully associate cue-outcome pairs, reported by predictive licking to rewarded cues

We trained mice in a Go/NoGo visual discrimination task to study reward association learning in a non-spatial context. We recorded the activity of single neurons from POR and MEC across the entire learning phase, from naive to expert performance. Food-restricted, head-fixed mice placed on a running wheel learned to associate drifting gratings of three different orientations (Plus Cue: 0°, Minus Cue: 270°, Neutral Cue: 135°) with rewarding, aversive or neutral cue outcomes depending on their choice of action (licking/no licking) following stimulus presentation. One training session contained 162-486 trials of cue-outcome pairing, where one trial consisted of a three-second stimulus window, followed by a two-second response window and a six-second inter-trial interval (Fig. 1a). Licking during the response window following the plus, minus, and neutral cues resulted in the delivery of a high-calorie milkshake (8-10 *µ*L Ensure), an aversive bitter solution (5 *µ*L of 0.1 *mM* quinine), or no outcome, respectively. To account for behavioral variability when comparing training sessions across mice, we defined the first session with a *d*^′^ *>* 1 (z-score of hit rate in all trials) as the learning session and the day with the highest behavioral score as the expert session. Typically, mice developed a pattern of predictive licking (i.e., licking during the stimulus window) in response to plus cues after successful cue-outcome pairing (example from one mouse in Fig. 1b). In expert sessions, animals began licking on average 0.79 ± 0.037 seconds earlier during plus trials compared to naive sessions, showing that the plus stimulus elicited predictive behavior long before reward delivery (Supplementary Fig. 1c). Mice typically reached expert performance, defined as a *d*^′^ of *>*2.0 within 5-10 days of training from the naive state (Fig. 1c). Based on hit rate (the proportion of trials with a lick response), animals appeared to recognize the reward-associated cue after three days of training, with hit rates approaching one by the expert stage. Interestingly, no clear distinction was observed between minus and neutral trials Fig. 1d and Supplementary Fig. 1e). Once mice achieved expert performance with the initial task rules, the cue-outcome pairing was reversed, assigning different outcomes to each visual cue (Plus Cue: 270°, Minus Cue: 135°, Neutral Cue: 0°).

**Figure 1:**
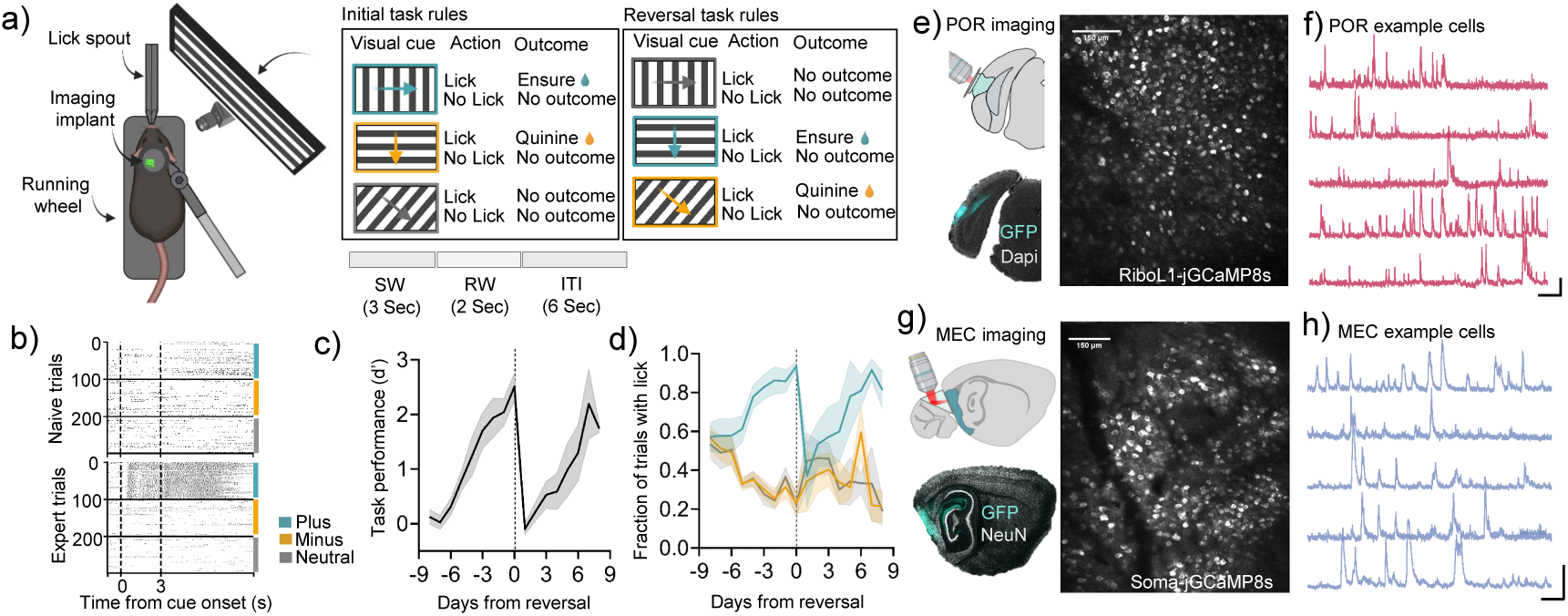
Behavioral performance and imaging while learning a visual-association task. a) Animals were head-fixed on a running wheel, and drifting gratings were displayed in their right visual field. Task rules before and after cue reversal are illustrated. b) Raster plot showing lick detection sorted by trial type (plus, minus, neutral) for one example animal in naïve and expert sessions. c) Behavioral performance score (d’) defined as (z-score of hit rate in all trials) across all animals and sessions, aligned to cue reversal (*N* = 9 pre, *N* = 4 post). * d) Fraction of trials with a lick response within the two-second response window after cue offset. * e) Coronal illustration of mouse brain showing the implanted optical window above postrhinal cortex (POR) (upper panel). The lower panel shows a representative example of the RiboL1-jGCaMP8s expression in POR. The right panel shows a two-photon imaging field of view from one example animal. The scale bar indicates 150 *µ*m. f) Δ*F/F*_0_ traces from five POR example neurons. The vertical scale bar indicates a 200% increase in Δ*F/F*_0_, and the horizontal scale bar indicates 15 seconds. g) Sagittal illustration of mouse brain indicating targeted Soma-jGCaMP8s expression in the MEC and position of the right-angle transcranial prism (2*x*2*mm*) (upper panel). The lower panel shows a representative example of the Soma-jGCaMP8s expression in MEC. The right panel shows a two-photon imaging example field of view from MEC. The scale bar indicates 150 *µ*m. h) Same as in f) for five MEC example neurons. *Data is shown as mean ± SEM across mice.

To record the activity of neurons in layer II/III of POR and dorsal MEC during learning, we used two-photon calcium imaging of locally expressed RiboL1-jGCaMP8s and Soma-jGCaMP8s, respectively. POR mice were implanted with a chronic optical window, and MEC mice were implanted with a right-angled transcranial prism (*N* = 4 mice from POR and *N* = 5 mice from MEC) (Fig. 1e-h shows imaging examples from POR and MEC). One day of recording consisted of a 15-minute baseline in darkness before 90 minutes of cue-outcome training, followed by 60 minutes in darkness after training.

### POR and MEC neurons develop response bias to rewarded trials

We first investigated whether neural activity in POR and MEC neurons reflected features of the Go/NoGo task across different stages of learning. In total, we recorded the activity of 46,181 neurons (non-unique) across 122 sessions in POR (25,491 cells over 54 sessions, averaging 472 cells per mouse per session) and MEC (20,690 cells over 68 sessions, averaging 304 cells per mouse per session). A substantial population of neurons in both regions exhibited changes in response profiles to one or more trial types (plus, minus, neutral) as training progressed (Fig. 2a, 150 top responsive neurons, sorted by maximum response and trial type from one POR and one MEC example mouse across naive and expert training stages before and after cue-outcome reversal). Neurons were classified as trial-active if they showed a significant increase in fluorescence in at least one 1-second time bin within the cue or response window relative to a 1-second baseline before cue onset. Consistent with previous findings, we observed that POR neurons were trial-active even during the naive training phase. Interestingly, MEC neurons were less responsive in the naive state. However, as learning progressed from naive to expert sessions, a significant fraction of neurons in both regions developed a response bias toward rewarded plus trials (POR: 27% ± 5% to 57% ± 10%, MEC: 17% ± 8% to 75% ± 4%) (Fig. 2b). Notably, we also observed neurons that displayed significant responses to minus (POR: 22% ± 8%, MEC: 9% ± 5%) and neutral (POR: 23%±8%, MEC: 14%±10%) trials (Fig. 2c). This trend was further supported by the trial-sorted mean population response during different training phases, both before and after cue reversal, which demonstrated that both POR and MEC populations developed a pronounced bias toward rewarded trials, as measured by the absolute mean population response in each region (Fig. 2d).

**Figure 2:**
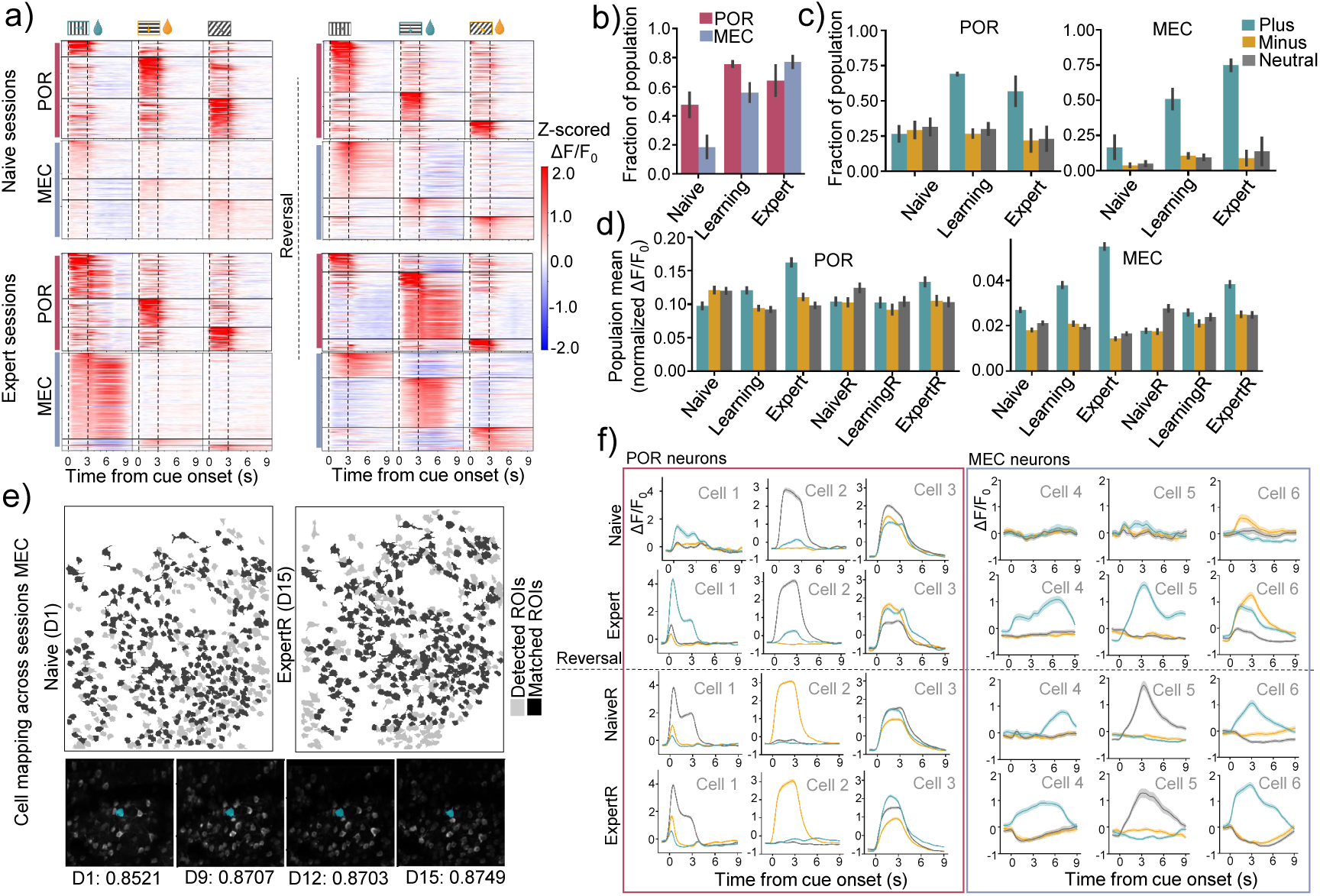
Neurons become modulated as learning progresses. a) Time course of mean activity (Z-scored Δ*F/F*_0_) in response to plus-, minus-, and neutral trials (columns) for the top 150 trial active neurons (rows) across different learning stages, before and after reversal, from one POR and one MEC example mouse. Cells are sorted by peak responses during the cue-or response-window. b) The trial active cell fraction increases with learning of the task. Trial active cells are defined as cells with a significant increase in Δ*F/F*_0_ for at least one time-bin (1s) within the cue-or response-window. * c) Same as b) but sorted by trial type across training phases. * d) Absolute mean population response (Δ*F/F*_0_ across stimulus window relative to one-second preceding stimulus onset) of all trial active neurons across all training stages in POR and MEC, before and after reversal. * e) Cell masks detected for individual imaging sessions (naïve and expert reversal) from the same animal in light grey, overlaid by cell masks of cells present in both sessions (dark grey). Example of a neuron (in blue) matched across multiple imaging sessions with an overlap score for each session (lower panel). f) Examples of single-cell activity. Mean activity time courses (Z-scored Δ*F/F*_0_) in response to the different trial types for six example neurons identified across sessions. *Data is shown as mean ± SEM across mice.

We tracked the same neurons across sessions to follow how individual neurons’ responses change during learning. At the start of each session, the field of view was consistently identified by referencing a selection of cells from the naive recording session. Detected ROIs were matched across sessions by alignment to other sessions using a transform between the field of views and computing the dice scores between pairs of cell masks (Fig. 2e, see Methods). We reliably tracked 296 ± 95 cells from POR and 316 ± 39 MEC from naive to expert (mean ± SEM across mice). At the level of individual neurons tracked across sessions, we observed that POR neurons showed stable tuning curves to specific cue orientations across reversal. In MEC, most trial active cells were biased to the rewarded trials across rule reversal, while a small fraction of neurons followed the cue identity (Fig. 2f). Interestingly, in MEC mice (example mouse shown inFig. 2a), the neuron population seemed to exhibit more robust responses to all cues after the reversal, including the cue that had never been rewarded.

### Learning enhances task-related population activity

Given that both regions underwent drastic changes in their responsiveness to plus trials as learning progressed, we employed a Generalized Linear Model (GLM) to investigate how the specific behavioral and task-related variables influence neural activity patterns, predicting the activity of individual neurons on a frame-by-frame basis across sessions (as in [Goltstein et al., 2021, Sugden et al., 2020, Steinmetz et al., 2019, Musall et al., 2019]; see Methods). The model included a total of eight regressor groups consisting of behavioral features (licking, speed, and brain motion), each of the three stimuli (plus, minus, neutral), and delivery of reward (Ensure) or the aversive quinine solution. Supplementary Fig. 3a illustrates an example design matrix encompassing all the regressors. The model was trained on a random selection of 70% of the trials in each session and evaluated on the remaining 30%. We validated the model’s performance using repeated random subsampling (averaged over 100 different train/test splits) and defined cells with 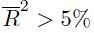 on the test sets as overall modulated.

Fig. 3a shows examples of predicted activity versus actual activity in two task-modulated cells from POR and MEC. This indicates the model successfully captures the main activity patterns in a subset of neurons from both regions. Overall, the proportion of task-modulated cells was higher in the POR (83%) compared to the MEC (36%) across all subjects and sessions, as well as for individual training phases (Fig. 3b). Additionally, the *R*^2^ scores for modulated cells were generally higher in POR than in the MEC (Fig. 3c), indicating that the model better explained neural activity in the POR. During learning, we observed a general increase in the model’s explanatory power in both regions, as indicated by higher *R*^2^ values (Fig. 3c), with significant distinction between the naive and expert stages (Fig. 3c), showing that task-related variables better explained the neural activity as learning progressed. Interestingly, the increase in the fraction of task-modulated neurons from naive to expert stages appeared to be more pronounced in the MEC (Fig. 3b).

**Figure 3:**
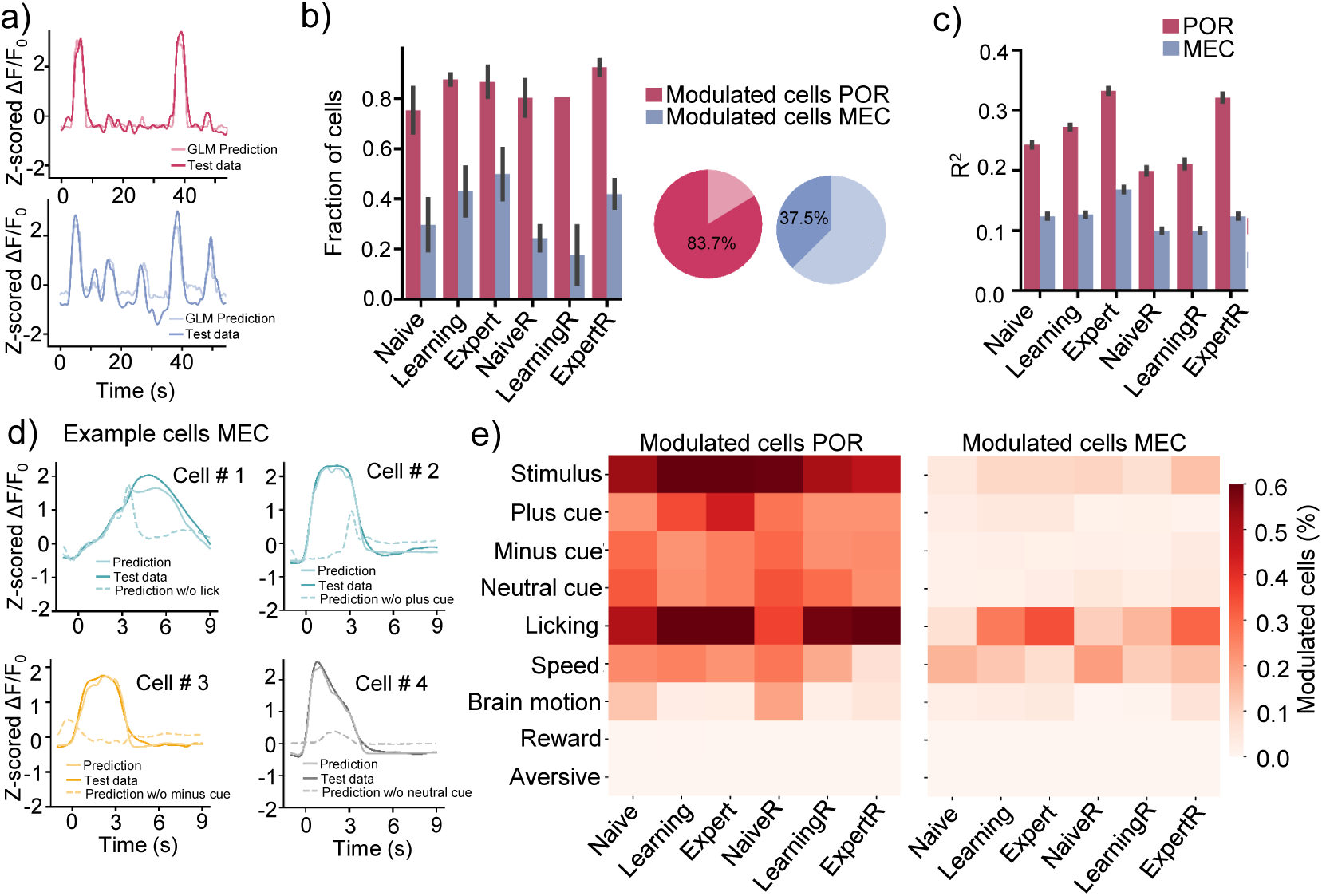
An increasing number of neurons becomes modulated by behavior– and task-related variables. a) GLM prediction versus actual neural activity for one example neuron from POR (upper panel, *R*^2^ = 0.927) and MEC (*R*^2^ = 0.678) in a 50s window. b) Fraction of neurons significantly modulated across the different learning stages for POR and MEC. Pie charts show the overall fraction of modulated cells in the respective regions. * c) Distribution of *R*^2^ values (full model including all regressors) across learning stages for POR and MEC. * d) Trial-averaged GLM prediction vs actual activity of MEC example cells. For each example cell, we show the model’s prediction, including all regressors, and the model without the regressor group, which has the highest uniquely explained variance. Only the most relevant trial type is shown. e) Heatmap showing the region-specific average fraction of modulated cells across all subjects (POR: *N* = 4 and MEC: *N* = 5) across different learning stages and regressors. *Data is shown as mean ± SEM across mice

To assess the individual contributions of each regressor group on modulated cells, we calculated the uniquely explained variance, 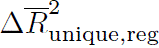, defined as the difference between the 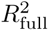 of the entire model and that of a model with the regressor in question shuffled (see Methods). A neuron was classified as modulated by a regressor group if the distribution of 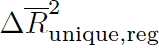 values significantly deviated from zero, and the unique contribution 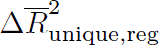 exceeded 2% (Supplementary Fig. 3d provides an overview of 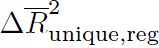, scores across different regressor groups), in trial-averaged predictions of neurons strongly modulated by different regressors. Models lacking the key regressor failed to capture significant aspects of the activity pattern, while the full models closely approximated the actual activity Figure 3d. The percentage of modulated cells across all subjects (POR: *N* = 4 and MEC: *N* = 5) at different learning stages is shown inFigure 3e. We noted an increase in lick-modulated cells from the naive to expert stages, both before and after the reversal stage (from 7% to 34% in MEC and 50% to 70% in POR before reversal) for both regions. Additionally, 61 ± 5% of neurons were modulated by the stimulus regressor group in POR across all stages, with fewer stimulus-modulated cells in the MEC (8 ± 2%).

### Subpopulations of neurons form clusters of task-relevant representations

While calculating the Δ*R*^2^ for each regressor group offers valuable insights into the individual contributions of each factor, it does not fully capture the complexity of neural tuning, particularly in cases where neurons exhibit mixed or temporally dynamic responses. To classify neurons based on their tuning profiles, we applied Agglomerative Clustering to the regression weights to identify clusters of cells with similar tuning characteristics. We first defined the licking, stimulus, reward, and aversive regressor groups as task-relevant and only used cells modulated by at least one of those groups. Overall, 81% of all recorded neurons in POR fulfilled this criterion, compared to 29% of the neurons in MEC. For the clustering process, each cell was represented by a vector containing all weights from the task-relevant regressors, excluding the intercept. By applying the GLM across sessions and animals, we could pool cells from all sessions and subjects. Importantly, clustering was based on the training sessions.

We utilized a consensus clustering scheme and found eight to be the optimal number of clusters (Supplementary Fig. 4a and b). We then removed two clusters due to their low number of cells (n¡ 50) from subsequent analysis (Supplementary Fig. 4c). Across all recording sessions, the clustering revealed two dominant clusters comprising a total of 10602 (40%) and 9685 (37%) cells, respectively, along with smaller clusters containing 2595 (10%), 1886 (7%), 1421 (5%), and 292 (1%) cells (Fig. 4a, relative numbers as a percentage of all relevant clusters C1-C6). We used UMAP [McInnes et al., 2020] to visualize the clusters in two dimensions, which showed that the clusters separated well, with the two dominant clusters positioned on opposite sides of the UMAP space (Fig. 4b). Analysis of the average weight vectors for the different clusters revealed that the dominant clusters, C1 and C2, were significantly influenced by the lick regressor, with C1 exhibiting a high negative weight and C2 a high positive weight associated with this regressor. Clusters C3-C5 were predominantly related to weights from one or more stimulus regressors (Fig. 4d-e).

**Figure 4:**
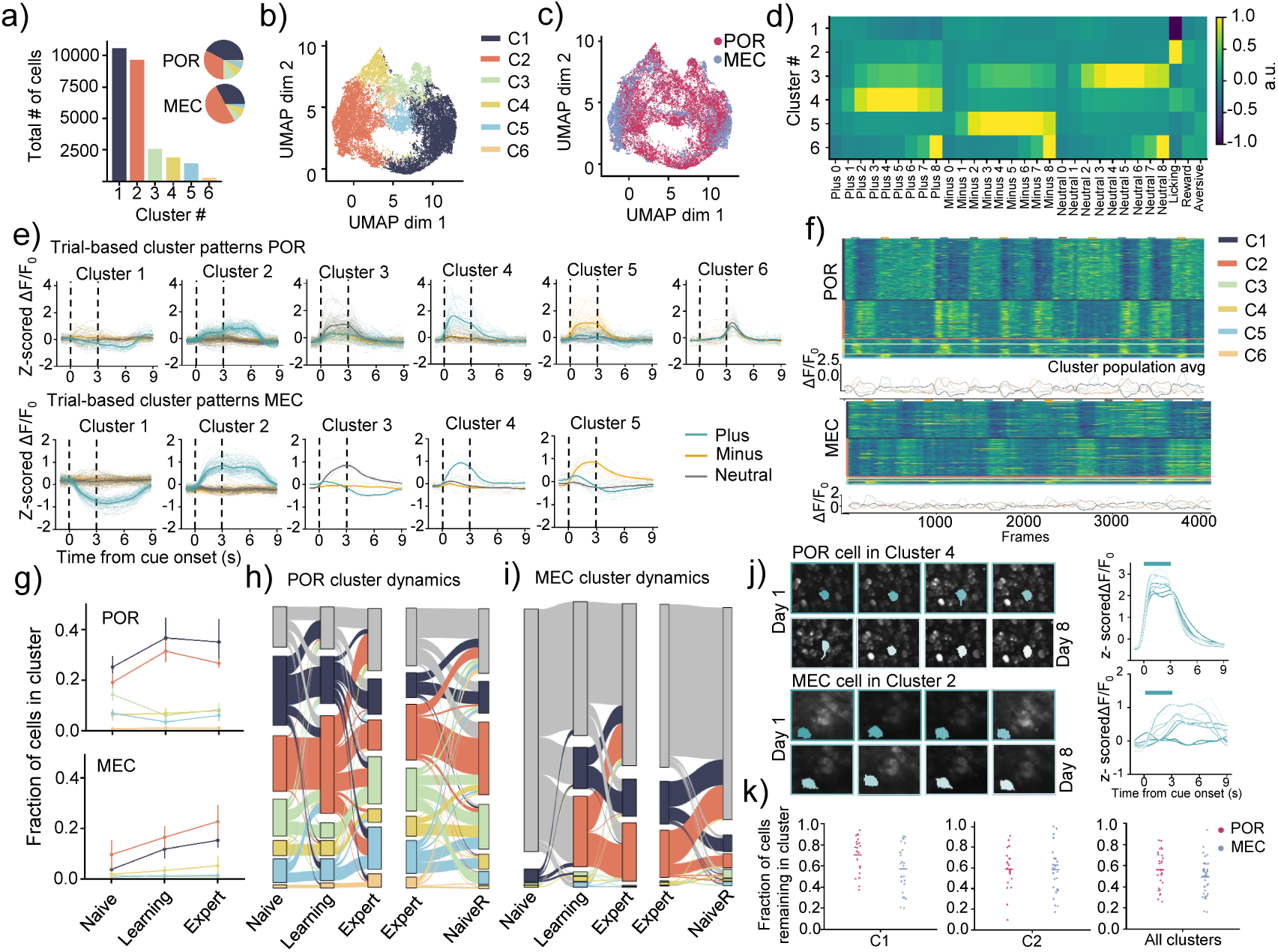
Clustering neurons with similar task-relevant tuning characteristics. a) Histogram of total cell counts in the seven identified clusters. The colored pie chart shows the fraction of each cluster per region (POR: postrhinal cortex, MEC: medial entorhinal cortex). b) Cells represented by their task-relevant regression weight vectors in two-dimensional UMAP space. c) Same as b) with region-specific coloring. d) The average weight vector of each cluster across sessions. Note the stimuli-induced activity in C3-C5. e) Trial-averaged responses of cells in each cluster for each trial type in an example session from POR (upper panel) and MEC. Thin lines indicate individual cell responses and bold lines indicate the respective mean of all cells in that cluster. C6 was not present in most MEC animals. f) Cell responses from an example session from POR (upper panel) and MEC (lower panel) are sorted by cluster, and each row represents one cell. Color bars on top indicate visual stimuli (plus cue: green; minus cue: orange; neutral: grey. Note the licking-related activity in C1 and C2 and the cue-modulated activity in C3-C5. g) Average fraction of cells in relevant clusters across different stages showing an increase for the two lick-related clusters C1 and C2 in both regions. h) Sankey diagram of cluster dynamics between learning stages for one POR example animal. i) Same as h) but for a MEC example animal. j) Examples of one POR and one MEC neuron matched across eight sessions before reversal. Left panels indicate two-photon FOVs highlighting with the cell mask consistently matched across sessions in shades of green. The right panel shows mean activity time courses (fractional change in fluorescence, Δ*F/F*_0_) for the same neuron across indicated sessions in response to plus trials. Both neurons remain within their respective clusters across all sessions. k) Cluster stability scores for pairs of subsequent sessions (each dot represents one such pair). The stability score measures the fraction of cells found in the same cluster in the subsequent session. Here, we only included C1 and C2 clusters and session-cluster pairs with stability higher than chance (see Methods).

The trial-averaged responses to different stimuli for cells in each cluster during a representative learning session for one POR and one MEC animal are shown in Fig. 4e. The influence of the lick regressor was particularly evident in C1 and C2, where C1 cells showed decreased activity and C2 cells showed increased activity during plus trials. This anti-correlation is even more apparent in the raster plot in Fig. 4f, where C2 cells were collectively active during plus trials, while the activity of C1 cells was suppressed. Additionally, increased stimulus-induced activity could be observed in C3-C5 during trials of the corresponding stimulus type. Cells in C6 responded relative to the offset of the stimuli (Fig. 4d and e) and were predominantly found in POR. Such offset-related activity is in line with a previous study [Bair et al., 2002].

### Behavior clusters were less apparent in MEC from the naive state

Throughout learning, the fraction of cells in C1 and C2 increased in both POR and MEC. In POR starting from higher baselines of 25 ± 4% (C1) and 19 ± 3% (C2) at the naive stage to 35 ± 9% (C1) and 27 ± 2% (C2) at the expert stage. In MEC from 4 ± 1% (C1) to 15 ± 3% (C1) and 10 ± 6% (C2) to 23 ± 7% (C2) (Fig. 4g; fractions of all cells). In contrast, the proportions of cells in clusters C3-C5 remained relatively stable across all stages, with percentages fluctuating within a consistent range (10 ± 2% C3, 7 ± 1% C4, 6 ± 1% C5 in POR, and 1 ± 0.3% C3, 4 ± 2% C4, and 1 ± 0.3% C5 in MEC). Generally, the stimulus-related clusters C3-C5 were bigger in POR than MEC, consistent with the higher percentage of stimulus-modulated cells described before.

For subjects with a stable field of view across all learning stages, we tracked the cluster membership of individual cells over time. Importantly, we could not follow all cells identified in each session. Hence, the subsets of neurons matched across sessions may not fully represent the cluster proportions in the overall population identified in any session. In MEC, we found that cells not initially modulated (grey) at the naive stage were recruited into clusters C1 and C2 during the learning. In contrast, many POR cells exhibited specific tuning from the naive session Fig. 4h and j. Although there were instances of cross-cluster dynamics and cells returning to the non-modulated group, many of the cells in C1 and C2 remained in their respective clusters once recruited (examples of cluster-stable cells from each region are shown in Fig. 4j). To quantify this stability, we calculated a cluster consistency score, which measures the proportion of cells in a cluster in one session that remains in the same cluster in the subsequent session.

We compared that to a shuffled distribution (see Methods). We only considered the C1 and C2 clusters due to the insufficient number of cells in the other clusters. Comparing cluster consistencies across regions revealed that clusters were significantly more stable than expected by chance (Supplementary Fig. 4h-k) and that POR showed tendencies of higher consistency across sessions compared to MEC (Fig. 4k).

In summary, clustering across regions revealed six task-relevant clusters, with a more significant fraction of POR neurons being task-modulated from the beginning (naive sessions). Compared to the MEC, the POR had more neurons representing both behavior and, especially, stimulus-related information. In contrast, the MEC exhibited a similar recruitment trend but only for behavior-related neurons, indicating different stimulus-encoding across regions.

### Stimulus clusters in POR encode more visual information

The fraction of stimulus-modulated cells (belonging to C3, C4, or C5) was higher in the POR compared to the MEC (Fig. 5a). To assess the visual information encoded by the different clusters, we applied a support vector machine (SVM) classifier, reporting the mean decoding accuracy of the trial type. We used a 5-fold cross-validation scheme, repeated ten times, based on the responses of all stimulus-modulated cells (C3-C5) from individual animals and training sessions. The overall decoding accuracy for these clusters was significantly higher in the POR compared to the MEC (Fig. 5b), which held true across different learning stages Fig. 5c) and specific stimuli (Fig. 5d). Notably, while decoding accuracy in the POR remained close to 1 across learning stages, the MEC showed increased accuracy for the plus stimulus as mice progressed from the naive to expert stages, indicating learning-induced stimulus encoding (Fig. 5d). Generally, the decoding accuracy in MEC was lower for fewer cells (Supplementary Fig. 5a), and most MEC sessions had fewer cells in those clusters, which could explain the lower decoding accuracy. However, even when comparing similar numbers of cells, decoding performance was typically better in the POR than in the MEC (Supplementary Fig. 5a).

**Figure 5:**
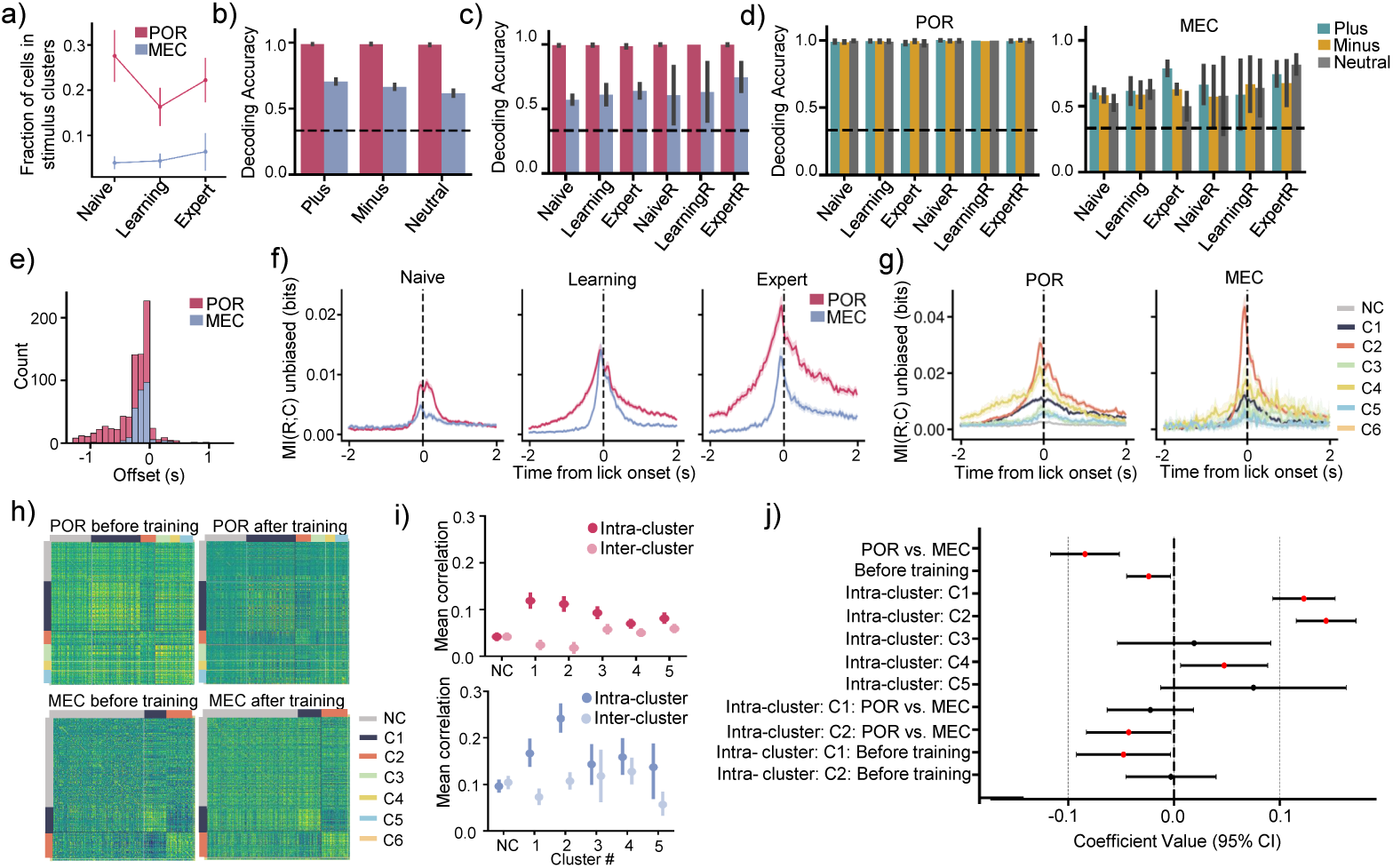
The POR encodes more visual information than the MEC, while both regions show increased choice information across learning with a more robust correlation structure MEC. a) Cell fractions in stimulus clusters across learning stages for the two regions. * b) Decoding accuracy for the stimulus clustered cells is higher in POR than MEC with no difference between plus-, minus-, and neutral cues. The dotted line indicates chance. * c) Region-specific decoding accuracy for stimulus clusters across all defined learning stages. * d) Region-specific and trial-type sorted decoding accuracy for stimulus clusters across learning stages. * e) Histogram showing the offsets between the lick signal and deconvolved traces for C2 cells (negative offset means spikes precede licking). f) Mutual information between neuronal response and choice (i.e., lick vs. no-lick, MI(R;C)), shown as the mean of the population. Overall, the choice information was virtually non-existent in naive mice but increased with learning in both brain areas. g) Same as in (f) but for each of the functional cell clusters. In both brain areas, the choice information was highest in clusters 2 and 4, i.e. those most closely linked to the rewarded cue-outcome pair (reward consumption and Plus-cue, respectively) h) Cluster-sorted correlations of a baseline run before training and a spontaneous run after training for one POR and one MEC animal. i) Distribution of intra-cluster versus inter-cluster correlations for the different clusters in POR (top) and MEC (bottom). j) Most relevant linear mixed effects model coefficients and their 95% confidence interval. Significant coefficients (*p <* 0.05) are marked in red (see Supplementary Fig. 5j for a complete list of coefficients). *Data is shown as mean ± SEM across mice ^∗^*P <* 0.05, ^∗∗^*P <* 0.01, ^∗∗∗^*P <* 0.001.

We further compared these results with two additional approaches: one using stimulus-modulated cells identified by GLM (excluding those modulated by running or licking) and another using only non-clustered cells (Supplementary Fig. 5b-c). While the GLM-based results were comparable, decoding accuracy was much lower when using non-clustered cells, even though more cells were used in the MEC sessions (Supplementary Fig. 5c). These findings indicate that the visual information decoded from the POR was significantly higher than from the MEC. Although MEC decoding levels were above chance, they reflected much lower levels of visual information compared to the POR. Interestingly, our results suggest a learning-related increase in stimulus encoding for the plus cue within the MEC.

### Information about behavioral choice develops with learning

Given that most task-responsive neurons were linked to reward consumption (C1 and C2), we examined this relationship more thoroughly. A key question was whether the activity of lick-modulated cells merely reflected the behavior or represented an internal process, with neural activity potentially preceding the behavior. To investigate this, we measured the temporal delay between licking and the activity of C2 neurons that showed elevated responses during lick behavior. Using deconvolved activity data, we calculated the conditional probability of a lick occurring, given the deconvolved signal at various time lags (see Methods). The lag with the highest conditional probability was defined as the temporal delay between neural signals and behavior. We compared these conditional probabilities to a null distribution and included only cells with probabilities above the 99th percentile of the shuffled distribution. This criterion was met by most (93%) cells in the C2 cluster (Fig. 5e). Results showed that, across subjects, the majority (93%) of this group of cells had a negative lag (−0.27 ± 0.01*s*), indicating that neural activity preceded licking behavior, suggesting it may reflect reward predictions or choice to respond.

To quantify how much information is encoded in the neuronal population about the behavioral decision, i.e. the decision to lick or not, we calculated the mutual information between the deconvolved activity trace of each neuron on a frame-by-frame basis relative to lick onset [Panniello et al., 2024]. We only included the first lick in bouts of at least 5 licks within 0.5 seconds, and only if the preceding 2 seconds did not include licks. We found that the mutual information between neuronal response and choice (i.e., lick vs. no-lick, MI(R;C)) increased with learning (Fig. 5f). Interestingly, when examining the choice information in each of the functional clusters, both C2 and C4 showed higher mutual information scores across sessions (Fig. 5g), supporting the notion that neurons in both of the reward-trial-related clusters encode information about the choice to lick.

### Behavior clusters identified during cue-outcome pairing exhibited stronger correlations during darkness runs in the MEC

Next, we investigated whether the cells within functional clusters would exhibit similar activity patterns during spontaneous activity before and after training. Previous studies, such as those involving grid cells, have shown that the activity of these cells correlates during periods of sleep [Gardner et al., 2019]. To test this, we compared the pairwise correlations of cells within clusters to those outside the clusters for baseline runs before training and spontaneous runs following training. A Linear Mixed Effects model accounted for inter-subject variability, day-to-day fluctuations, and missing data points. In this model, region, cluster identity, time point (pre– and post-training), and correlation type (intra-cluster versus inter-cluster) were treated as fixed effects, with all interaction terms included (see Methods). Subject identity and session were modeled as random effects.

For clusters C1 and C2, within-cluster correlations (intra) were significantly higher than inter-cluster correlations in the non-clustered cells (Fig. 5i and j) across MEC and POR regions. However, this difference was more pronounced in the MEC than in the POR (Fig. 5j). Additionally, while clusters C3-C5 tended to exhibit higher intra-cluster correlations, this was statistically significant only for C4 and with a smaller effect size than C1 and C2. Examples of pairwise correlation structures that illustrate this effect is shown in Fig. 5j. The model further revealed that pairwise correlations were generally higher in the MEC than in the POR and increased after training. These findings suggest that the identified functional clusters form intrinsic groups that maintain consistent activity patterns outside the task context in which the clustering analysis was performed initially.

### Emerging functional clusters also cluster anatomically

Previous studies have found that grid cells in MEC also form anatomical clusters [Obenhaus et al., 2022, Gu et al., 2018, Heys et al., 2014, Dombeck et al., 2009]. Therefore, an interesting question is whether the functional clusters we identified also show topographic organization in the tissue. To answer this question, we first plotted the cell masks of the two big clusters, C1 and C2, and found that cells in MEC tend to cluster anatomically (Fig. 6a). We then used a subsampled k-nearest neighbors approach (similar to [Obenhaus et al., 2022]) to determine the distances within clusters versus a shuffled distribution of randomly assigned cluster identities (see Methods). We required at least 12 cells to be in a cluster in a particular session to get stable results for the analysis. Only clusters C1 and C2 had sufficient numbers of cells in enough sessions to be included in this analysis.

**Figure 6:**
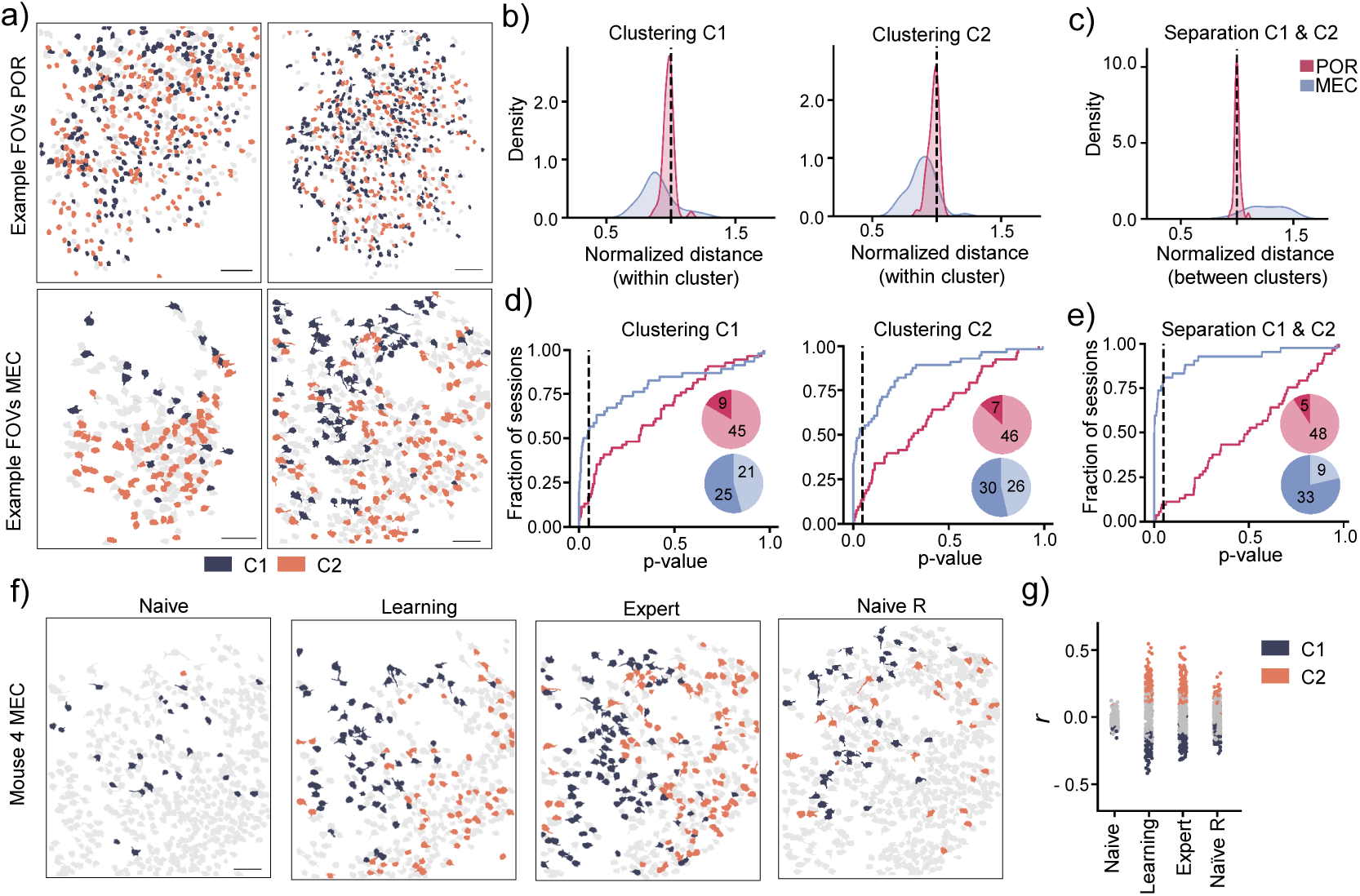
Functional cell groups are also anatomically clustered in MEC. a) Anatomical distribution of cell masks of C1 (blue) and C2 cells (orange) in two example mice from each region. All other cells are shown in light grey. The scale bar indicates 150 *µ*m. b) Distributions of subsampled k-nearest neighbor (*k* = 5) distances within the clusters C1 and C2 compared to a baseline with shuffled cluster identities in POR and MEC. c) The Distribution between C1 and C2 distances to a shuffled distribution shows that the clusters are also significantly separated in MEC. d) Fraction of eligible sessions (more than 12 cells in each cluster) across all mice that show significant anatomical clustering of C1 cells (left panel) and C2 cells (right panel). e) Fraction of sessions with significant separation between C1 and C2 cells across regions. f) Anatomical distribution of cell masks for C1 and C2 cells in a representative MEC mouse at different learning stages (from naive to naive reversal sessions). Note the recruitment of task-modulated cells from naive to expert and the anatomical clustering. The scale bar indicates 150 *µ*m. g) Distribution of R-values for the lick regressor across all detected cells in different sessions throughout the learning stages for the mouse shown in panel f. Cells belonging to clusters C1 and C2 are color-coded accordingly.

If a functional cluster also clusters anatomically, one would expect the distribution of distances to be lower than the shuffled distribution (Fig. 6b). Indeed, we found that in most sessions (54%), C1 and C2 were significantly clustered in MEC. In POR, however, we observed clustering in fewer sessions (C1: 17% of sessions and C2: 13% of sessions) (Fig. 6b and d). We then investigated whether the clusters would be spatially segregated in anatomical space. One possibility is that both functional clusters could occupy the same region in the field of view. By comparing intercluster distances with those from shuffled cluster identities Fig. 6c and e), we found that in most MEC sessions (79%), the clusters were significantly separated, while in POR, only 10% of sessions showed significant separation. This indicates that functional clustering corresponds to topographic organization in MEC but not POR. Notably, in the MEC and POR, the recruitment of cells to these clusters was learning-dependent Fig. 6f-g and supplementary Fig. 6e). However, anatomical clustering and separation were apparent in the MEC whenever the functional clusters were present.

## 3 Discussion

We have demonstrated that neurons in the dorsal MEC undergo significant changes in their response profiles as mice learn non-spatial visual associations. A substantial portion of MEC neurons developed a response bias toward rewarded trials, modulated not by the visual stimulus but by the behavioral approach and consumptive behavior. In contrast, a larger fraction of neurons in POR were activated by the visual stimuli before the animals learned the task. Furthermore, we show that MEC neurons show anatomical clustering that is correlated with their functional properties.

Although the MEC receives substantial visual input, MEC neurons did not show much response to visual stimuli in naive animals. In contrast, a significant fraction of POR neurons exhibited selective visual tuning to all three cue orientations in the naive state with a developing population bias to the rewarded cue upon successful cue-outcome pairing, in line with previous studies using the same behavior task [Ramesh et al., 2018, Sugden et al., 2020]. The amount of visual information that could be decoded from POR neurons was substantially higher than the MEC, where visual decoding accuracy was slightly above chance levels (Fig. 5b-d). While others have shown that MEC neurons integrate visual stimuli in virtual navigation tasks [Kinkhabwala et al., 2020, Nguyen et al., 2024, Campbell and Giocomo, 2018], although with less visual drive compared to primary visual and retrosplenial cortex [Campbell et al., 2021], we observed only sparse visual representations, even in well-trained animals. However, the reversal data suggest that the initial encoding of cue orientation is retained across the learning of both rule sets, indicating modest levels of visual information also in the MEC.

However, from the representations of stimuli, it is not evident that the MEC simply inherits the visual information from the POR. Although the source of this visual information is not explored in this paper, recent studies indicate that the POR primarily projects to the lateral entorhinal cortex [Doan et al., 2019] and that the MEC receives visual input from multiple sources beyond the POR [Shao et al., 2024, Olsen et al., 2017, Dubanet and Higley, 2024]. Additionally, integrating environmental cues is crucial for correcting errors during spatial navigation [Hardcastle et al., 2015, Evans et al., 2016]. Thus, the sparse visual representations observed in this study contrast with the considerable representation of visual cues reported by others [Kinkhabwala et al., 2020] and might hint at the idea that some spatial requirements gates significant integration of visual information in the MEC. It is also plausible that spatially modulated cells, such as grid cells, and non-spatially modulated cells are differentially influenced by sensory input Casali et al. [2019], Kinkhabwala et al. [2020], Clark and Nolan [2024]. Furthermore, the structured representations of sensory information in the entorhinal cortex, both auditory [Aronov et al., 2017] and visual [Wilming et al., 2018]) in tasks not requiring navigation in physical space suggests that this hypothetical gating mechanism may also apply to sensory processing within cognitive space. However, such integration may not occur when the sensory information lacks structure.

Particularly in the MEC, we found that behavior-related clusters exhibited correlation structures that may suggest action coding, as they maintained similar correlation patterns during darkness runs. However, unlike reports of non-specific action coding across the brain [Steinmetz et al., 2019], the behavior-correlated neurons in this study appear to represent task-specific information. The precision in neuronal correlates paralleled the development of the behavior, with higher precision when the animals learned the tasks, and thereby, the behavior was goal-oriented and accurate. Upon cue reversal, when behavioral performance was significantly impaired, neural responses changed, showing a loss of cell members in the functional behavior-related clusters and weaker correlations. This could also be interpreted as encoding choice information [Panniello et al., 2024], as supported by our observation that neural activity preceded the onset of behavior (see Fig. 5). Indeed, when we computed the mutual information between neuronal activity and behavioral choice, we found that this increased with learning in both brain areas, an increase that was most profound in the reward-related clusters that encode reward consumption and plus-cue visual stimulus. These findings suggest that neuronal activity patterns are conditional, predicting reward-conditioned movements [Lee et al., 2020].

In perceptive decision-making, goal-approach behavior likely follows cognitive processes that are closely associated with instructed but also uninstructed movements [Stringer et al., 2019, Niell and Stryker, 2010, Saleem et al., 2013, Musall et al., 2019, Cohen and Kohn, 2011]. Thus, a key question is whether cognitive processes can be distinguished from their associated movements. This can be solved experimentally, e.i. by training animals to withhold the behavioral response during stimulus presentation, ensuring separation of perceptive decisions and action [Panniello et al., 2024]. In this study, we attempted to deal with it from an analysis point of view and leave movement-related regressors out to assess their individual contributions [Musall et al., 2019]. However, this approach assumes that cognitive and movement-related processes are distinguishable and, thus, likely driven by separate neural processes. Alternatively, cognitive processes like perceptive choice and movement could be closely connected [Cisek and Pastor-Bernier, 2014, Cisek and Kalaska, 2010], and might require specific approaches for proper evaluation [Hasnain et al., 2024]. Here, we demonstrate that neurons in the behavior-related clusters exhibit changes aligned with task performance. Notably, the representation of licking behavior in these clusters during expert sessions is markedly different from that in the naive state, despite unconditioned licking behavior across all trials in the naive state, both before and after the reversal (Fig. 4).

Our findings reveal a striking anatomical organization within the MEC population, in which the correlation structure of the network is reflected in the anatomical organization in the tissue (Fig. 6). This is in line with reports on the clustering of functionally related groups of neurons in the MEC [Obenhaus et al., 2022, Gu et al., 2018, Heys et al., 2014, Dombeck et al., 2009]. Typically, in the brain, the anatomical organization of neurons mirrors the organization of their input, as seen in sensory and motor systems (e.g., retinotopy in POR [Wang and Burkhalter, 2007]). If this principle can be applied to the organization of correlating cells in this study, it is conceivable that the POR and MEC either receive input from different sources or that the intrinsic structure of the MEC network drives this type of organization [Pastoll et al., 2013, Fuchs et al., 2016, Couey et al., 2013], or a combination of both.

Although identifying the input that drives the observed neural responses in MEC is beyond the scope of this paper, the observed neural correlates closely parallel the response profiles observed in the basolateral amygdala (BLA) and VTA-originating BLA-terminating axons in the same task [Lutas et al., 2019]. This could suggest that the gradual changes in neural correlations of reward-related behavior observed in this study may result from reinforcing signals to the cue-reward-evoked engagement of the midbrain dopaminergic system. [van Zessen et al., 2021, Saunders et al., 2018, Lee et al., 2021, 2020, Tye et al., 2010, Esber et al., 2012, Bissière et al., 2003]. Here, the dopaminergic neurons shift their responses from reward-responsive to also responding to reward-predicting cues Schultz et al. [1997], Day et al. [2007]. Interestingly, the dopaminergic drive of the BLA has been linked to unavoidable aversive [Fadok et al., 2009, Lutas et al., 2019], and appetitive associations, but not to avoidable mildly aversive outcomes (e.g., bitter tastants) [Lutas et al., 2019], which might explain the lack of clear separation between the negative and neutral trials observed in our study.

Future studies should address the relevance of these MEC computations for behavioral performance, aiming to disentangle cognitive processes from associated movements, possibly through experimental designs that allow for the separation of visual stimuli, choice, and approach behavior [Hasnain et al., 2024]. Further work is needed to understand the projections’ nature and function from POR and BLA to the MEC. Moreover, given the unresolved problem of self-motion versus environmental sensory input to the MEC, a speculative hypothesis for future investigation is that the transition from a non-spatial to a spatial context will shift some gating mechanism, possibly reflecting shifts in brain state to alter the influence of the visual system on the MEC.

## 4 Methods

### 4.1 Animals

All experiments were conducted at the animal facility at the Department of Biosciences, Oslo, Norway, according to the Norwegian Animal Welfare Act and the European Directive 2010/63/EU, after approval by the National Animal Research Authority of Norway (Mattilsynet, FOTS 14680 and 29491).

Adult C57/BL6j mice (*>* 12 weeks old) from Janvier Labs were used for MEC experiments, while PV-Cre mice (strain # 017320, RRID: IMSR JAX:017320) mice were used for POR experiments. The data collected from PV-Cre mice, used as a comparison group in this study, also served as a virus control group in Lensjø et al., 2024 (unpublished). In these animals, RiboL1-jGCaMP8s was co-injected with AAV-PHP.eb-hsyn-DIO-mRuby3 to achieve PV+ interneuron-specific expression. The experimental protocol in Lensjø et al., 2024 (unpublished) included bi-daily injections of clozapine N-oxide (CNO) following training sessions. Spontaneous recordings from those days were excluded from the analysis in this study. All mice were housed in GM500 IVC cages for at least two weeks before beginning any procedures. After surgery, the mice were housed individually for the entire experiment. The cages were maintained in a housing room with a light intensity of 215 lux on a 12/12-hour light cycle. The room temperature was 21 ± 1°C, with 25-30% humidity. For enrichment, each cage contained a plastic tunnel and ample nesting material. The mice had ad libitum access to food and water unless otherwise specified.

### 4.2 Surgical procedures

The animals were anesthetized using isoflurane (3% induction at 1L/min and 1-2% maintenance at 70 mL/min), followed by subcutaneous injection of buprenorphine (0.05 mg/kg, Indivior Ltd) for general analgesia and bupivacaine (Aspen) into the scalp for local analgesia. An intramuscular injection of dexamethasone (5 mg/kg, MSD Animal Health) was administered to prevent edema. The skin was cleaned with 70% ethanol and Jodopax before the skin covering the top of the skull was removed. The periosteum and other membranes were carefully cleared using fine forceps and cotton swabs.

### 4.3 Virus injection and optical implants

A 3 mm circular craniotomy was performed at coordinates –4.65 AP and 4.35 ML for virus delivery to the POR. A handheld Perfecta 300 drill (W&H) with a 0.5 mm bit (Hager & Meisinger GmbH) was used, ensuring the surrounding bone was thinned for uniformity. Glass capillaries, beveled at a 40-degree angle, were prepared and loaded with hSyn-RiboL1-jGCaMP8s PHP.eB virus using a Nanoject 3 (Drummond Scientific, USA). The capillary was inserted 0.4 mm deep, and 300 nL of viral solution was injected in 5 nL steps. For the MEC, a small craniotomy was created 3.3 mm lateral to the midline, anterior to the transverse sinus, and posterior to the lambdoid suture. The pipette was filled with 200 nL of hSyn-Soma-jGCaMP8s PHP.eB virus and inserted at a 5-degree angle. Injections were administered at depths of 1.45 mm and 1.65 mm, with the pipette held in place for 10 minutes after injection. A custom titanium head post was affixed to the skull using cyanoacrylate, stabilized with VetBond (3M) and C&B Metabond (Parkell), aligned with the skull plane. For the POR, a 3 mm cranial window (Tower Optical) was attached to a 5.0 mm cover glass (Warner Instruments) using UV-cured Norland Optical adhesive (Thorlabs GmbH, Germany) and positioned flush with the skull. For the MEC, a 2×2 mm craniotomy was made 2.7-4.7 mm lateral to the midline, with edges posterior to the lambdoid suture and anterior to the transverse sinus. The dura mater was incised along the transverse fissure, and a right-angled prism (Tower Optical) was bonded to a 5.0 mm cover glass, then lowered into the fissure until the cover glass was flush with the skull. The implant was secured with C&B Metabond, and a 3D-printed light shield was attached to the head post using black dental acrylic. Post-surgery, mice received subcutaneous injections of 0.5 mL saline and meloxicam (5 mg/kg, Boehringer Ingelheim VetMedica GmbH), with meloxicam treatment continuing for three days.

### 4.4 Habituation and visual association task

Training sessions began 2-3 weeks after surgery, depending on the expression levels of Soma-jGCaMP8s. Before imaging experiments, the animals were food-restricted to 85% of their ad libitum weight and habituated to head fixation for three days. During habituation, the mice were head-fixed on a custom 3D-printed running wheel using optical posts mounted to the optical table with holding clamps (Standa). On days 1-2 of habituation, the mice were introduced to liquid food (Ensure) delivered through a handheld syringe. On day three, the mice were habituated to a custom 3D-printed capacitance-sensing lick spout with one tube for Ensure delivery and one for quinine, placed in front of their mouths so that the tongue of the mouse touched both ports upon licking. For habituation, Ensure was delivered in 8 − 10 *µ*L aliquots through the lick spout upon licking. Following habituation, the mice underwent intermittent Pavlovian plus cue training and non-Pavlovian plus cue trials (go/lick trials) for 10-20 minutes on two consecutive days to associate the onset of cues on the monitor with reward deliveries and action requirement (licking), respectively.

After pre-training, experimental imaging sessions commenced. Each session consisted of a 15-minute baseline run in darkness, followed by 30-90 minutes of cue-outcome trials, with the duration determined by the animals’ motivational state. Food-restricted mice were trained to discriminate square-wave drifting gratings of different orientations (2 Hz and 0.04 cycles/degree) generated using the open-source Python software PsychoPy37 and synchronized with two-photon imaging through a parallel port and PCIe-6321 data acquisition board (National Instruments). A 3-second stimulus window displaying either the plus cue (0°), minus cue (270°), or neutral cue (135°) on a monitor covering the visual field of the right eye was followed by a 2-second response window, which, depending on the action of the mouse (lick/no-lick), triggered Ensure, quinine, or no outcome, respectively. After the response window, a 6-second inter-trial interval separated the previous trial from the next. Licking outside the response window had no outcome. The order of trial types was pseudorandom, with equal numbers of the different trial types in each session. Finally, a 60-minute recording of spontaneous neuronal activity in darkness was conducted, during which the animals could run on the wheel. Mice were considered proficient in the task when their performance exceeded a *d*^′^ value of 2 (Stanislaw and Todorov, 1999). Following successful cue-outcome pairing (*d*^′^ *>* 2), the cues were reversed, meaning each cue was assigned a new outcome. For the reversal, task structure and training sessions were performed as previously described, but the different outcomes had to be associated with new cue orientations: plus (270°), minus (135°), and neutral (0°) cues. The described method was adapted from [Burgess et al., 2016] and [Ramesh et al., 2018].

### 4.5 Wide-field and two-photon imaging

To assess the quality of the prism implant, expression levels, and anatomical targeting of Soma-jGCaMP8s, single wide-field images were acquired by a Canon EOS 4000D camera through a ×5 Mitutoyo long working distance objective (0.14 NA) in an Olympus BX-2 microscope. The light source was a xenon arc Lambda XL lamp (Sutter Instruments) with 480/545 nm filters (# 39002, Chroma). The mice ran freely on the running wheel during imaging. For in vivo two-photon imaging, we used a resonant-galvo Movable Objective Microscope (MOM, Sutter Instruments) with a MaiTai DeepSee laser (SpectraPhysics) set to a wavelength of 920nm. The objective was a Nikon ×16 objective (NA 0.8), giving a field of view of approximately 1050×890µm. The laser was controlled by a Pockel’s cell (302 RM, Conoptics), and fluorescence was detected through Chroma bandpass filters (HQ535-50-2p and HQ610-75-2p, Chroma) by PMTs (H10770PA-40, Hamamatsu). Images were acquired at 30.9Hz using MCS software (Sutter Instruments). Output power at the front aperture of the objective was measured with a FieldMate power meter (Coherent) and set to 45-55mW, depending on the viral expression. The microscope was tilted to an angle of 10–20 degrees during imaging depending on the surface angle of the prisms, in addition to the 4-6 degree forward tilt resulting from the head post rotation. Reference images were acquired before the first imaging session and used to identify the same field of view in consecutive imaging sessions.

### 4.6 Immunohistochemistry

An intraperitoneal injection of ZRF-cocktail (zolazepam; 18.7 mg/ml, tiletamine; 18.7 mg/ml, xylazine; 0.45 mg/ml, fentanyl; 2.6 µg/ml, 0.75 ml/g body weight) was administered before transcardial perfusion with PBS followed by 4% paraformaldehyde in PBS. Brains were dissected and post-fixed for 24 hours at 4°C. Following cryoprotection in 30% sucrose in PBS, brains were flash frozen, and 40 µm sagittal sections were made using a cryostat (Leica). Sections were rinsed thrice in PBS, followed by 1-hour blocking in 2% bovine serum albumin in 0.3% Triton-X-100 in PBS. Primary antibodies were diluted 1:1000, and free-floating incubation was conducted at 4°C with constant agitation. Sections were rinsed three times in PBS and incubated with secondary antibodies diluted 1:500 in PBS, followed by additional rinsing in PBS. Sections stained with Neurotrace Nissl (NeuroTrace 640/660 Nissl, Invitrogen) were diluted 1:100 in PBS, followed by 10 minutes in 0.1% Triton-X-100 in PBS after primary and secondary incubation steps. The antibodies used were chicken anti-GFP (RRID: AB 2534023, Invitrogen, 1:1000 dilution, rabbit anti-NeuN (RRID: AB 2532109, Abcam), goat anti-chicken AlexaFluor 488 (RRID: AB 142924 Life, 1:500 dilution) and goat anti-rabbit AlexaFluor 647 (RRID: AB 2535813). Finally, sections were rinsed in PBS, mounted on Superfrost Plus adhesion slides, washed in dH2O, and coverslipped with mounting medium (Ibdi). Tile scans were used to acquire overview images of brain sections on an Andor Dragonfly spinning-disc microscope equipped with a motorized platform. Stitching with 20% overlap was performed using Fusion software (Bitplane). The Andor Dragonfly was built on a Nikon TiE inverted microscope with a Nikon PlanApo ×10/0.45 NA objective.

### 4.7 Image analysis

Motion correction and automatic detection of regions of interest (ROIs) were performed using Suite2p [Pachitariu et al., 2016]. All detected cell masks were manually curated based on bright-ness, cell mask shape, and signal-to-noise ratio. To calculate relative fluorescence Δ*F/F*_0_, traces were first corrected for neuropil contamination using *F_c_* = *F* − 0.7 ∗ *F*_neu_ + 0.7 ∗ *F*_neu,median_, where *F* represents the raw fluorescence, *F*_neu_ is the neuropil signal defined by Suite2p as the fluorescence in a surrounding region around the ROI, and *F*_neu,median_ is the median value of that signal (adapted from [Goltstein et al., 2018]). Δ*F/F*_0_ was then calculated as (*F_c_* − *F*_0_)*/F*_0_, where *F*_0_ is the running median (with a window size of 600s) of the neuropil-corrected trace *F_c_*. The OASIS algorithm was used to deconvolve the Δ*F/F*_0_ traces [Friedrich et al., 2017].

### 4.8 Cell matching across sessions

We developed a custom script to create pairwise mappings between the ROI IDs of sessions captured from the same FOV. To map the cells of two sessions, we began by calculating a similarity transform between the mean images of the FOVs of the recordings, incorporating rotation, translation, and uniform scaling [Reddy and Chatterji, 1996]. This transform was then applied to the binary cell masks of the second session, followed by re-binarization of the masks using a threshold of 0.2. To identify potential cell pairs, we calculated the Dice similarity coefficients between the cell masks of the two sessions. A cell from the first session was mapped to the cell in the second session with the highest Dice score, provided that the score exceeded 0.5 and the gap to the next highest candidate was at least 0.15. These criteria allowed us to establish a mapping table between the two sessions.

Mapping of the various sessions began by selecting one session as the reference, and then all subsequent sessions were mapped to the reference. After completing this process, we selected the next session as the reference and repeated the mapping for all other sessions, excluding those previously used as references. During this iterative mapping process, many cells were mapped indirectly (e.g., if two cells from different sessions were mapped to the same cell in the reference session, they were considered mapped to each other). We focused our efforts on mapping cells that had not yet been matched. Ultimately, we compiled a coherent mapping table that included all sessions for a given subject from the same FOV. To ensure the accuracy of our mappings, we generated cell maps for all groups. We performed a manual quality check, excluding cells from the mapping if they did not exhibit sufficient similarity.

### 4.9 GLM analysis

To assess the individual effects of behavioral and task variables on neural activity, we employed a generalized linear model (GLM) to predict the frame-wise activity of individual neurons [Goltstein et al., 2021, Sugden et al., 2020, Steinmetz et al., 2019, Musall et al., 2019]. Each stimulus was encoded by shifted Gaussian basis functions (*σ* = 0.25*s*) with time shifts of −0.5*s*, 0*s*, 0.5*s*, 1.0*s*, 1.5*s*, 2.0*s*, 2.5*s*, 3.0*s*, and 3.5*s* relative to stimulus onset. Note that each stimulus remained on the screen for 3 seconds. The basis functions spanned the entire on-screen duration, with additional time before and after stimulus presentation to account for potential pre– and post-stimulus effects. Additionally, a lick regressor capturing all lick events and a running regressor using the instantaneous running speed were included. We also incorporated a brain motion regressor based on the frame-by-frame movement of the imaging frame, which was calculated as the Euclidean distance in pixels. All regressors were smoothed using a Gaussian filter with *σ* = 0.25*s*. Finally, we added two regressors: one aligned to reward delivery and the other to aversive delivery, represented by Gaussian basis functions centered at the respective frames (*σ* = 0.25*s*). Regressor groups were then defined based on one or more related regressors: one group for each stimulus type (containing all shifted basis functions for that stimulus), a general stimulus regressor group comprising all stimulus-related regressors, and groups for licking, running, brain motion, reward, and aversive events, each containing the relevant regressor. The dependent variable was the Gaussian-filtered (*σ* = 0.25*s*) and Z-scored Δ*F/F*_0_ signal of each neuron.

We used an elastic net regularization with a combined L1 and L2 penalty. We trained and evaluated the model on a subset of sessions (expert sessions of all animals) to identify suitable model parameters. The parameter set that yielded the highest median *R*^2^ across all neurons was selected (Supplementary Fig. 3b). The chosen parameters were *α* = 0.001 and l1-ratio = 0.9, which were applied to all subsequent analyses.

The general training and evaluation procedure used repeated random subsampling with 100 iterations. In each iteration, we split the data randomly into 70% training and 30% testing data based on trials. The model was fit to the training set and evaluated on the test set using the *R*^2^ score for each neuron. To assess the model’s robustness, we randomly shifted all individual regressors in time (between 0.05*T*_session_ and 0.95*T*_session_, where *T*_session_ is the session duration) and re-evaluated the model using the same trials. This process generated distributions of 100 unshuffled 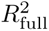 and 100 shuffled 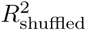 scores for each neuron in each session. To quantify the unique contribution of each regressor group, we further calculated the variance that was uniquely explained for each group. This was done by randomly shifting all regressors within a group in time (while maintaining the temporal structure between regressors within a group), then re-fitting and evaluating the model, similar to [Goltstein et al., 2021]. This yielded 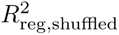 scores for each group across iterations. The uniquely explained variance of a regressor group was calculated as follows:

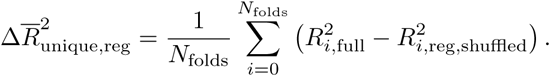

This value represents the variance explained exclusively by that regressor group and serves as a measure of its importance.

We considered a neuron overall modulated if the full model explained a significant amount of variance (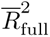). The threshold was set using the 95th percentile of the 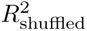 values when all regressors were shuffled, which gave a value of 4% (Supplementary Fig. 3c). Based on that, a threshold of 5% was chosen. To determine whether an individual regressor group modulated a neuron, two criteria were applied: (1) the uniquely explained variance of the regressor had to be at least 2%, and (2) a Wilcoxon signed-rank test was performed to compare the distributions of 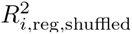 and 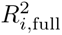 scores for each neuron. We applied Benjamini-Hochberg FDR correction for multiple comparisons, with a significance set at *p <* 0.01.

### 4.10 Timing between licking and neural activity

To determine the temporal lag between licking and neural activity, we first binarized the deconvolved traces of cells by setting time points with deconvolved activity greater than 0.01 to 1 and all other points to 0 to remove noise (following Panniello et al. [2024] and). For each cell, we computed the conditional probability *P* (*r*|*s*) between its binarized deconvolved activity *r* and the binary lick signal *s* at different lags using:

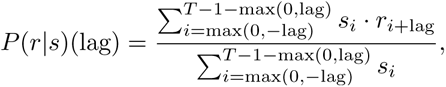

For a particular lag, where *T* is the number of frames. Lags ranged from –40 frames (approximately 1.3s) to 40 frames. We identified the lag that produced the highest conditional probability and compared it to a shuffled distribution where the lick signal was randomly shifted in time. This process was repeated 100 times, and we required the original probability to exceed the 99th percentile of the shuffled distribution. This analysis was restricted to cells whose activity increased in response to licking, specifically C2 cells. Of all C2 cells, 92.51% met the percentile criterion, and for these, we report the lags that maximized the conditional probability.

### 4.11 Clustering procedure

To identify groups of task-active cells with similar tuning properties across sessions and subjects, we clustered the regression weights obtained from the GLM analysis. Task-relevant regressor groups were defined as licking, individual stimuli, combined stimuli, reward, and aversive. We selected only those cells modulated by at least one task-relevant regressor group based on the modulation criteria outlined earlier. For each selected cell, an average weight vector was computed by averaging the regression weights of all task-relevant regressors over the 100 training iterations, excluding the intercept (as previously described). Clustering was then performed using a consensus clustering procedure, following [Monti et al., 2003] with Agglomerative clustering of these weight vectors, applying average linkage as the criterion and correlation as the similarity measure.

The consensus clustering procedure involved subsampling 80% of the data across 100 iterations for cluster numbers ranging from 2 to 20. We generated a consensus matrix for each cluster number that quantified how often two neurons were assigned to the same cluster relative to how frequently they were subsampled together. Ideally, stable clustering would result in consensus values close to either 0 (indicating two neurons never clustered together) or 1 (indicating two neurons always clustered together). To evaluate clustering robustness, we first computed the empirical cumulative distribution function (eCDF) for each consensus matrix and then calculated the area under the curve (AUC) of the eCDF. A high AUC indicates that most values are near 0 or 1, reflecting stable clustering, while a lower AUC suggests less stable clustering with many values between 0 and 1. We plotted the AUC values and selected eight as the optimal number of clusters using the Elbow method, where the AUC showed minimal improvement (≤ 0.01) for cluster numbers greater than 8 (Supplementary Fig. 4a and b). Finally, we applied Agglomerative clustering on the consensus matrix (which served as the similarity measure) with *n* = 8 to obtain the final clusters. For subsequent analyses, we excluded clusters 7 and 8, which were stable but included less than 50 cells (Supplementary Fig. 4c).

### 4.12 Decoding

We performed decoding using three different groups of cells. The first group was stimulus cluster cells, which included all cells from clusters C3, C4, and C5, identified as stimulus-related and present in all subjects. Second, we selected cells based directly on the GLM results. These cells were modulated by the combined stimulus regressor groups (or one of the individual stimulus regressors) but not by running or licking, as we aimed to isolate stimulus-related activity without behavioral confounders. The third group consisted of cells not included in the clustering analysis, meaning they were not modulated by any task-relevant regressor group (supplementary Fig. 5b and c).

For each cell group, we created a dataset by averaging the Gaussian-filtered (*σ* = 0.25*s*) and Z-scored Δ*F/F*_0_ signal over the 3-second stimulus window of each trial. Each stimulus type was treated as a distinct class, and each trial was represented by the population vector of the selected cells. We then applied a support vector machine (SVM) classifier with a linear kernel, using a 5-fold cross-validation scheme repeated ten times. Decoding accuracy was reported as the mean accuracy across all repetitions.

### 4.13 Mutual information

The mutual information between neural activity and choice was computed similar to what has been in done in Panniello et al. [2024]. However, in contrast to Panniello et al., we used sequences of licks as decision points. In brief, we first identified time points at which the subject licked at least 5 times in a 0.5s window and with no licks in the preceding 2s. We used the first lick onset of the sequence as the center and extracted windows of –2 to +2s around that center. We then randomly collected the same amount of sequences from periods with no licking (and at least 2s from any lick). The lick sequences were encoded with 1 and the no lick sequences with 0. Deconvolved traces were binarized using a threshold of 0.01 similar to Panniello et al. [2024] and Runyan et al. [2017] as described above (see Section 4.10). We then computed the cell-wise mutual information like Panniello et al. using

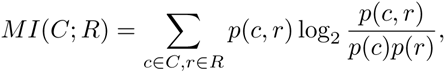

where *c* denotes the choice (0 or 1), and *r* the activity (0 or 1). Finally, we applied the Panzeri-Treves bias correction method to correct for limited sampling biases [Panzeri and Treves, 1996].

### 4.14 Linear mixed effects model

We employed a linear mixed effects model to investigate the intra-and inter-cluster correlation structure before and after training. This model accounted for repeated measurements and the variability arising from measuring different subjects on different days ([Lindstrom and Bates, 1988]). Random effects were modeled as random intercepts at the subject level, using subject ID as the grouping variable to account for individual-level variability. Additionally, day-level variation was included in the model as a random variance component. The fixed effects in the model were cluster ID (NC, C1, C2, C3, C4, C5, C6), correlation type (intra-or inter-cluster), timepoint (before or after training), and region (POR and MEC), with all interaction terms included. The model was optimized using the Newton-Raphson solving algorithm to ensure robust parameter estimation.

### 4.15 Anatomical clustering and separation

To investigate whether the identified functional clusters exhibited anatomical clustering or separation, we employed a k-nearest neighbor approach inspired by [Obenhaus et al., 2022]. For anatomical clustering, we required that each cluster contain at least 12 cells within a session. We then calculated the mean k-nearest neighbor distances across cells within that cluster for different values of *k* (ranging from 2 to 9). To generate a null distribution, we randomly selected a size-matched group of cells from the overall population and computed the mean distances to the k-nearest neighbors within this random group. This process was repeated 1000 times, and we calculated the p-value as the percentile of the actual distance in the shuffled distribution. A functional cluster was considered anatomically clustered in a session if *p <* 0.05. For the main analysis, we used *k*= 5, although we demonstrate in Supplementary Fig. 6 b and c that the choice of *k* has little impact on the results.

A similar approach was used to assess the anatomical separation between two clusters. In this case, both clusters were required to contain at least 12 cells. We then computed the mean k-nearest neighbor distances from cells in cluster A to cells in cluster B and vice versa and averaged the two values. To generate a null distribution, we performed 1000 iterations, where we randomly selected a size-matched cluster B’ and computed the mean k-nearest neighbor distances from A to B,’ followed by choosing a size-matched cluster A’ and calculating the same distances from B to A.’ The mean of these values was computed for each iteration, and the p-value was calculated as one percentual distance in the shuffled distribution. Cluster pairs with *p <* 0.05 were considered anatomically separated. To plot the distances, we normalized them by dividing them by the mean of the shuffled distributions.

## 5 Data availability

The original raw data from the imaging experiments will be deposited in a public repository in the future and will be available upon request from the authors.

## Acknowledgments

We thank Arvind Kumar for his insightful contributions to the analysis and to Guro Vatne for her contributions to the production of jGCaMP8s-carrying viruses. We are also grateful to Alessio Buccino for his assistance with the initial behavioral setup and to Constantin Bechkov for his helpful contributions to the analysis and input on the manuscript. This work was supported by the Research Council of Norway (Grants No. 248828, 250259, and 309547 to MF and AMS and grant no. 325892 to TH) and the University of Oslo’s Strategic Research Initiative CINPLA. This project has received funding from the European Union’s Horizon 2020 research and innovation programme under the Marie Skl-odowska-Curie grant agreement N° 945371.

## 7 Supplementary Information

The supplementary information contains supplementary figures S1–6.

**Supplementary Figure 1.**
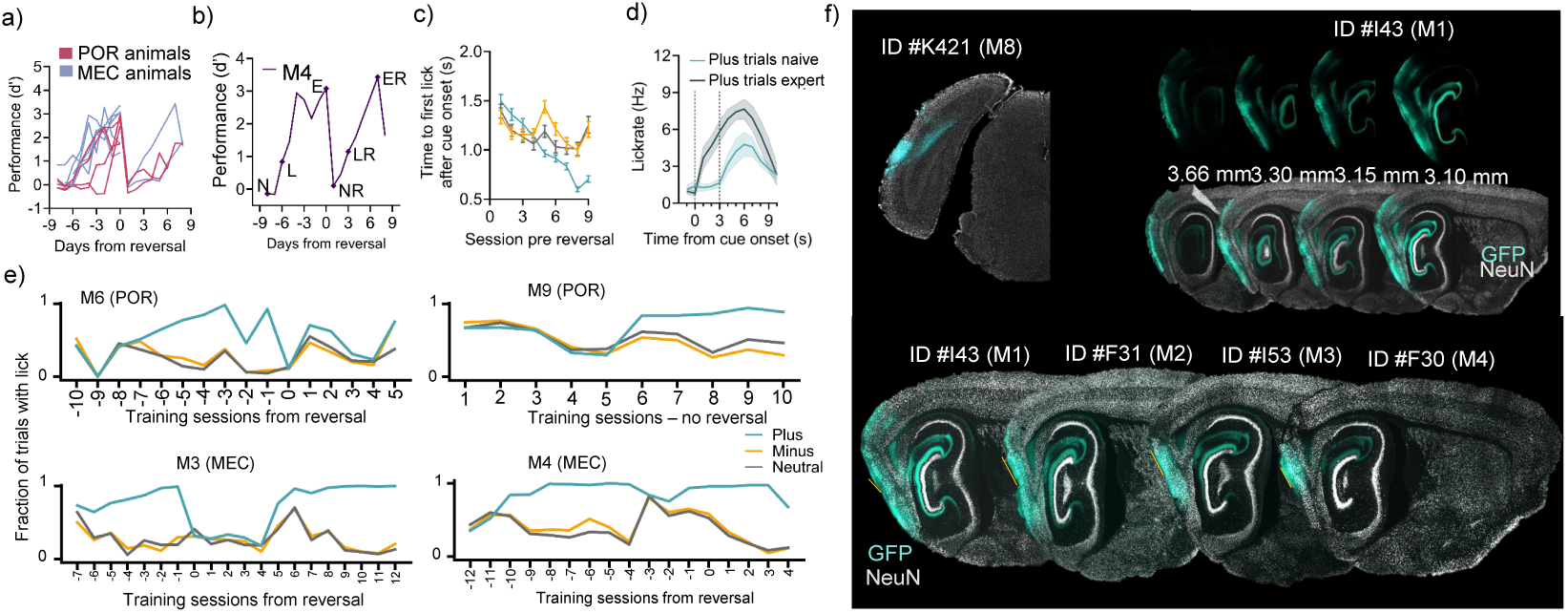
a) Behavioral performance scores (z score of hit rate and false alarm rate) for individual POR and MEC animals, aligned to cue reversal (*N* = 4 POR animals and *N*= 5 MEC animals). b) Performance score for mouse 4 (MEC animal) with naive (N), learning (L), expert (E), naive reversal (NR), learning reversal (LR), and expert reversal (ER) sessions indicated. c) Trial sorted time to first lick after cue-onset across animals from naive to expert stages pre-reversal. * d) Lick rate to plus trials within the trial structure for naive (light green) and expert (dark green) sessions. * e) Fraction of trials with a lick response within the two-second response window after cue offset for two POR and two MEC example animals. f) Coronal sections from one POR animal indicating RiboL1jGCaMP8s expression in the imaged hemisphere (upper left panel). Upper right panel shows a series of saggital sections from one MEC-targeted mouse (M1) indicating the spread of Soma-jGCaMP8s expression from lateral (3.66mm) to more medial coordinates (3.10mm). Saggital sections indicating the expression center of Soma-jGCaMP8s targeted to MEC from four mice. Orange bar indicates visible impact on the tissue from the transcranial prism (2*x*2*mm*) implant (lower panel). *Data is shown as mean ± SEM across mice.

**Supplementary Figure 2.**
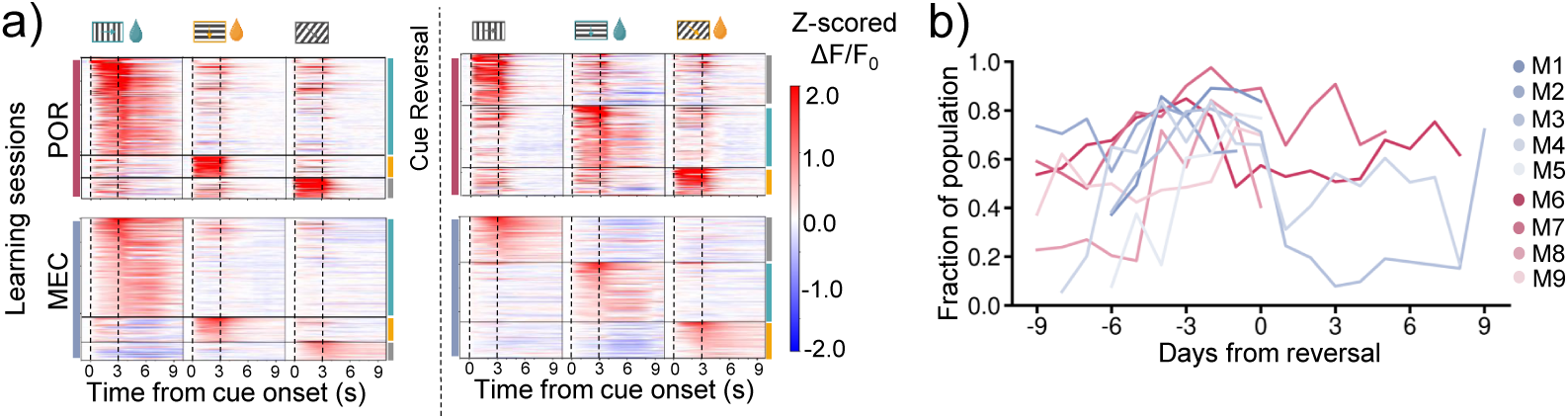
a) Time course of mean activity (Z-scored Δ*F/F*_0_) in response to plus-, minus-, and neutral trials (columns) for the top 150 trial active neurons (rows) in the learning session, before and after reversal, from the same POR and MEC example mice shown in Fig. 2a. Cells are sorted by peak responses during the cue-or response-window. b) The total fraction of trial active cells for individual animals. Trial active cells defined as cells with significant increase in Δ*F/F*_0_ for at least one time-bin (1s) within the cue-or response-window.

**Supplementary Figure 3.**
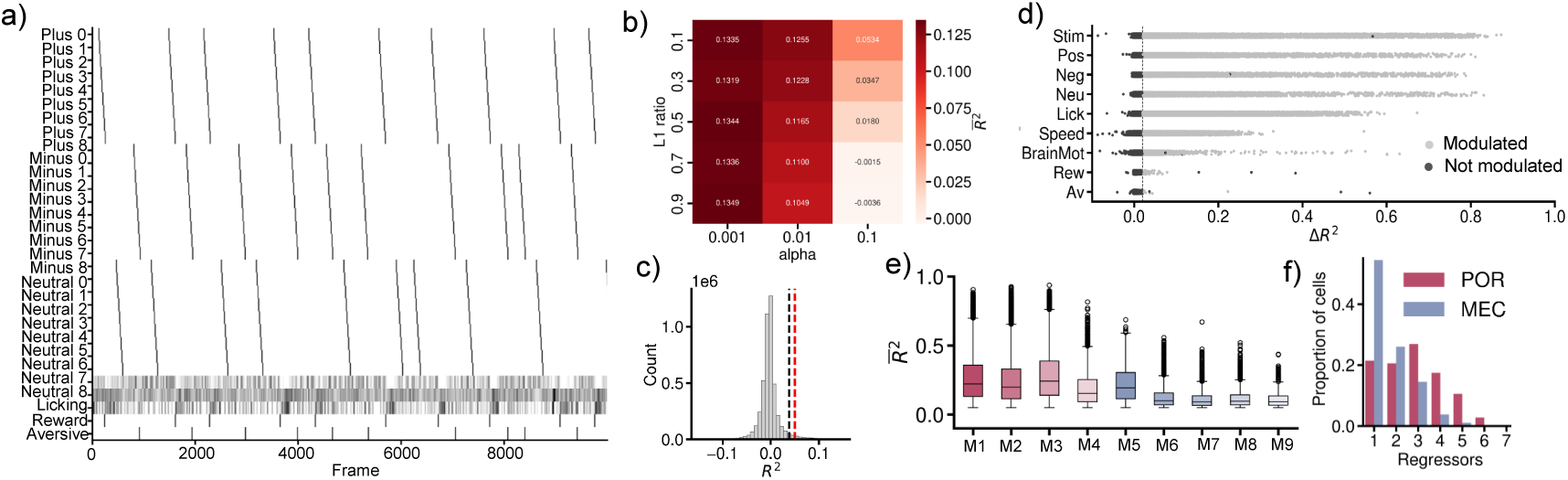
a) Example design-matrix including all regressors for the Generalized Linear Model (GLM). b) Median *R*^2^ values of the initial parameter grid search using expert sessions. The parameters leading to the highest value (*α* = 0.001 and l1-ratio = 0.9) were used for all subsequent experiments. c) Distribution of all *R*^2^ values of models where all regressors were shuffled. The black line indicates 95*th* percentile and red line indicates chosen threshold at *R*^2^ = 5% d) Distribution of 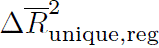 values for the different regressors across all sessions for POR and MEC. Cells modulated by the different regressors are indicated in light grey. e) Distribution of *R*^2^ values of modulated cells for individual animals across sessions. f) Fraction of cells modulated by single or multiple regressors across regions.

**Supplementary Figure 4.**
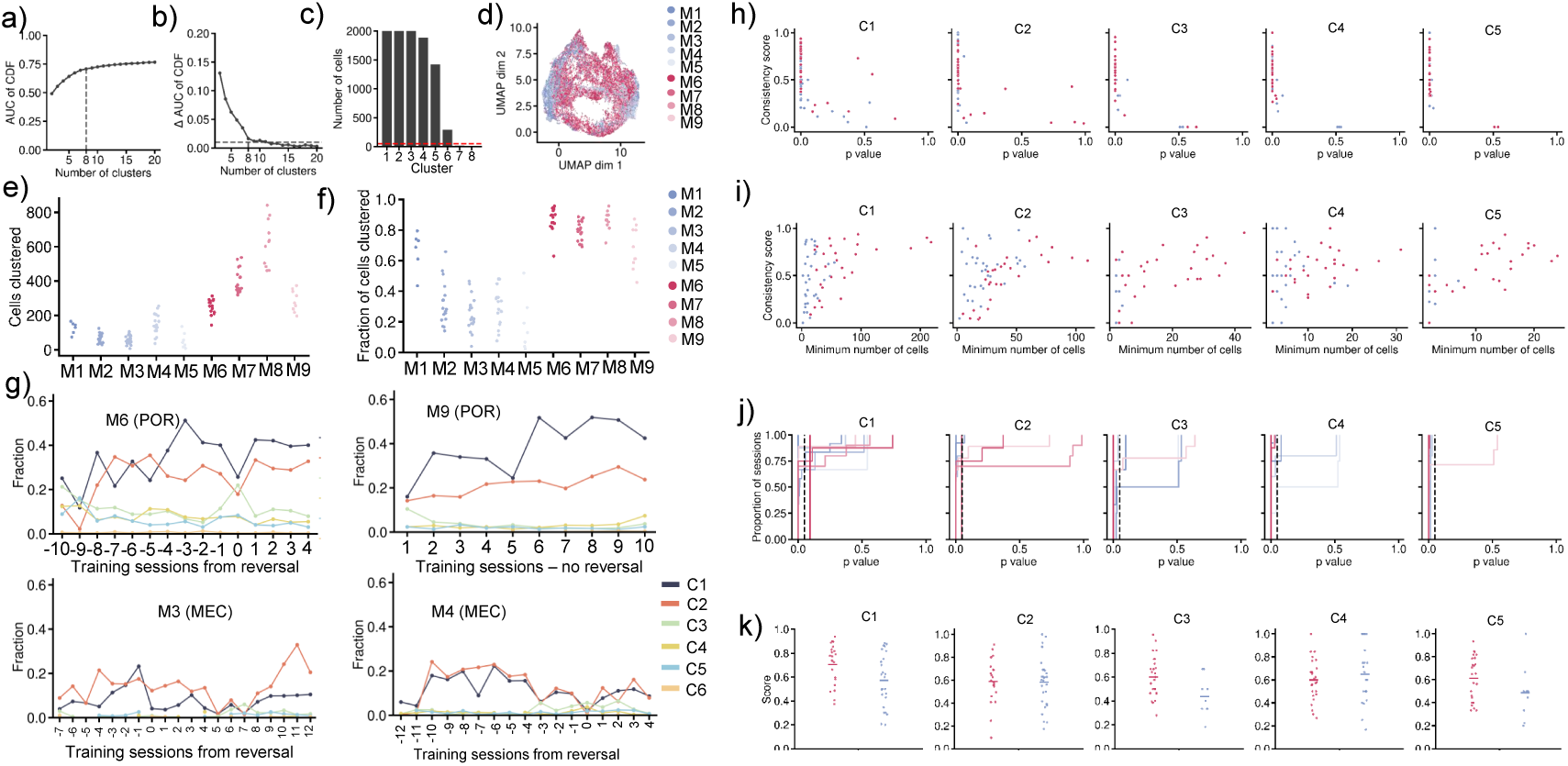
a) Area under the curve (AUC) of the eCDF calculated for each consensus matrix. b) Eight clusters were selected as the optimal number of clusters using the Elbow method, where the AUC showed minimal improvement (*≤* 0.1) for cluster numbers greater than eight. c) Histogram showing the total number of cells in each cluster. We set the inclusion criterion for further analysis to a minimum of 50 cells per cluster. d) Cells represented by their task-relevant regression weight vectors in two-dimensional UMAP space color-coded by region-specific animal ID. e) Total number of cells clustered per session for the nine different animals. f) Same as e) but showing the total fraction of cells clustered. g) Fraction of cells in C1-C6 for two POR and two MEC example animals across sessions. h) Relationship between cluster consistency scores and the p-values determined using a shuffled distribution (see Methods). Each dot represents one session. i) Relationship between consistency scores and the minimum number of cells of a cluster in the two relevant sessions used to determine the score. j) eCDF plots of the sessions of each animal over the p-value determined for the consistency scores of each session-cluster pair. k) Distributions of significant consistency scores for all clusters. Each dot represents one session (see Methods).

**Supplementary Figure 5.**
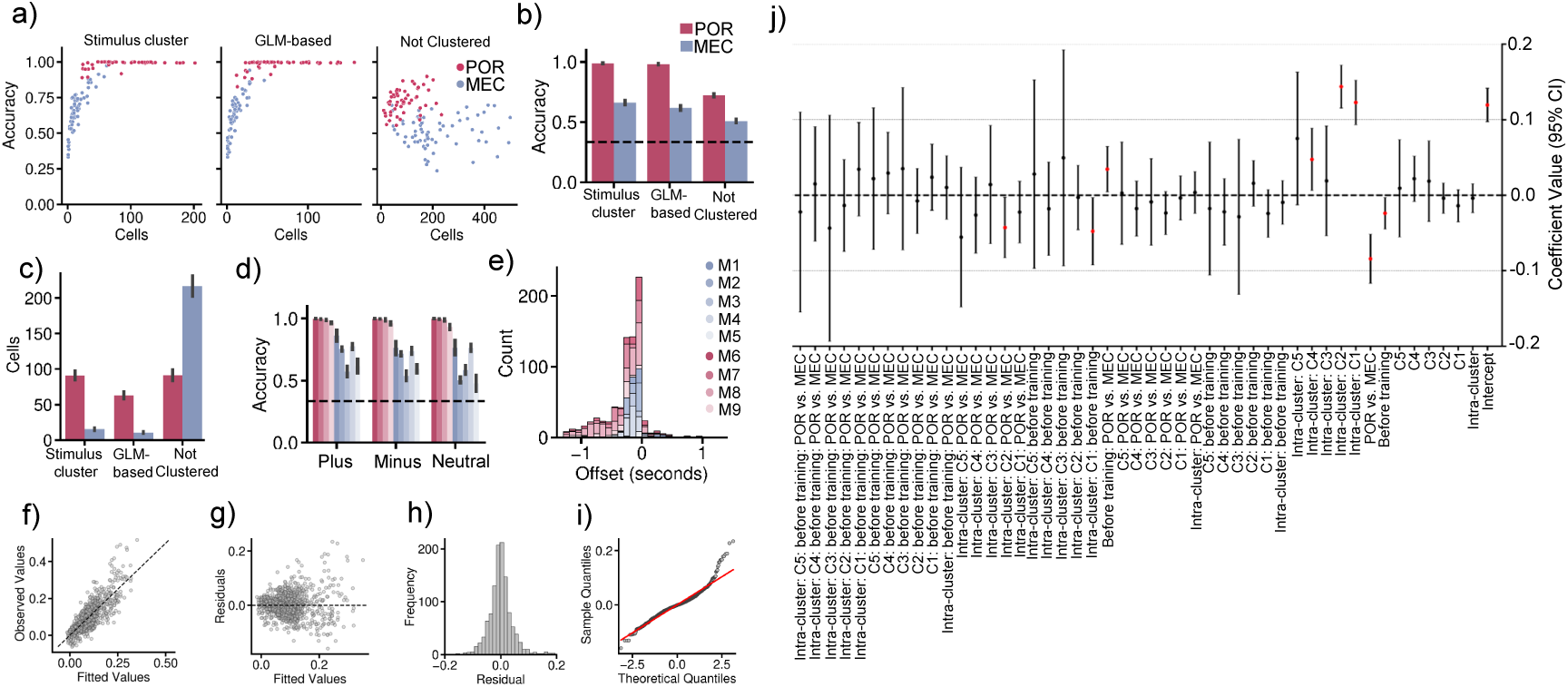
a) Mean decoding accuracy per cell using a support vector machine (SVM) classifier with a linear kernel, using a 5-fold cross-validation scheme repeated 10 times. Data points are color-coded according to the respective regions. b) Overall decoding accuracy for the three different cell groups tested: stimulus clusters (C3-C5), GLM-based (stimulus modulation determined by GLM), and not clustered cells. c) same as b) reporting absolute number of cells per region, per cell group. d) Decoding accuracy for the stimulus cluster, sorted by stimulus type for individual animals, color-coded by animal ID. e) Histogram showing the distribution of offsets between the lick signal and deconvolved traces for modulated cells in individual animals (negative offset means spikes precede licking). f) Scatterplot of the fitted values vs observed values in the linear mixed effects model applied to the pairwise correlations. g) Residuals over fitted values of the linear mixed effects model did not show a systematic trend. h) Distribution of residuals. i) Q-Q plot of residuals of the linear mixed effects model. j) All coefficients of the linear mixed effects models and their 95% confidence intervals. All significant coefficients (*p <* 0.05) are marked in red.

**Supplementary Figure 6.**
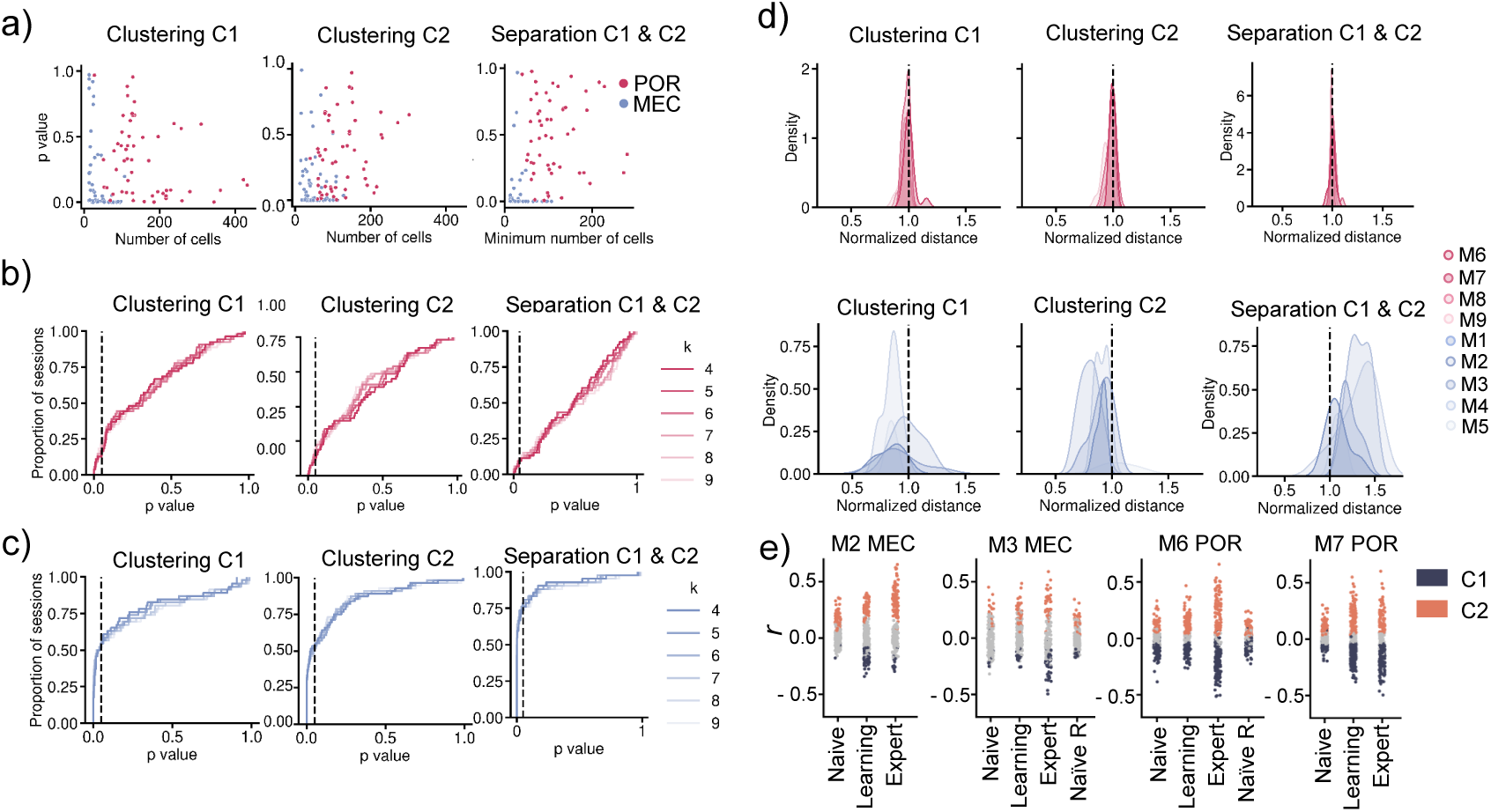
a) Relationship between p-values and number of cells for the clustering of C1 (left), and C2 (center), and the separation of the two (right). Each dot represents a session. For the separation, the minimum number of cells of C1 and C2 was chosen. b) eCDF plots for the p-values of the clustering of C1 and C2 and the separation of the two for different values of *k* in POR. c) Same as in b) but for MEC sessions. d) Distributions of normalized mean distances (normalized by the shuffled distribution) C1 cells (left) and C2 cells (center) as well as the distances between C1 and C2 (right) for different animals.

